# *Chlamydomonas reinhardtii* formin FOR1 and profilin PRF1 are optimized for acute rapid actin filament assembly

**DOI:** 10.1101/096008

**Authors:** Jenna R. Christensen, Evan W. Craig, Michael J. Glista, David M. Mueller, Yujie Li, Jennifer A. Sees, Shengping Huang, Laurens J. Mets, David R. Kovar, Prachee Avasthi

## Abstract

The regulated assembly of multiple filamentous actin (F-actin) networks from an actin monomer pool is important for a variety of cellular processes. *Chlamydomonas reinhardtii* is a unicellular green alga expressing a conventional and divergent actin that is an emerging system for investigating the complex regulation of actin polymerization. One actin network that contains exclusively conventional F-actin in *Chlamydomonas* is the fertilization tubule, a mating structure at the apical cell surface in gametes. In addition to two actin genes, *Chlamydomonas* expresses a profilin (PRF1) and four formin genes (FOR1-4), one of which (FOR1) we have characterized for the first time. We found that unlike typical profilins, PRF1 prevents unwanted actin assembly by strongly inhibiting both F-actin nucleation and barbed end elongation at equimolar concentrations to actin. However, FOR1 stimulates the assembly of rapidly elongating actin filaments from PRF1-bound actin. PRF1 further favors FOR1-mediated actin assembly by potently inhibiting Arp2/3 complex-mediated actin assembly. Furthermore, *for1* and *prf1-1* mutants, as well as the small molecule formin inhibitor SMIFH2, prevent fertilization tubule formation in gametes, suggesting that polymerization of F-actin for fertilization tubule formation is a primary function of FOR1. Together, these findings indicate that FOR1 and PRF1 cooperate to selectively and rapidly assemble F-actin at the right time and place.

**SUMMARY STATEMENT:** The *Chlamydomonas reinhardtii* formin FOR1 initiates rapid assembly of fertilization tubule actin filaments from monomers associated with the actin-assembly inhibitor profilin PRF1.

## INTRODUCTION

The actin cytoskeleton is a dynamic system important for diverse cellular processes. *Chlamydomonas reinhardtii* expresses a single conventional actin, IDA5, with 90% identity to mammalian actin, as well as an unconventional actin, NAP1 (Kato-Minoura, 1998; Lee et al., 1997), with low identity to mammalian actin (64%). *ida5* mutants have limited phenotypic consequences (Kato-Minoura et al., 1997), likely because NAP1 is upregulated upon IDA5 perturbation (Hirono et al., 2003, Onishi et al., 2018) and has compensatory functions (Jack et al., 2019). The presence of a perinuclear F-actin network has been recently established by both a fluorescent Lifeact peptide (Avasthi et al., 2014, Onishi et al., 2016) and an optimized phalloidin labeling protocol (Craig and Avasthi, 2019, Craig et al., 2019). Additionally, a population of F-actin also localizes at the base of the flagella, where it is important for flagellar assembly and proper intraflagellar transport (Avasthi et al., 2014, Jack et al., 2019). Thus far, cytokinesis has not been shown to require an F-actin network, as proliferation is Latrunculin B (LatB) and Cytochalasin D insensitive and *ida5* null mutants produce normal cleavage furrows (Onishi et al, 2016, Harper et al., 1992, Kato-minoura et al., 1997). Cytokinesis may instead utilize LatB-insensitive NAP1, which is upregulated during LatB treatment (Onishi et al., 2016, Onishi et al., 2018). Alternatively, a different mechanism of cytokinesis may be used in these cells.

Another clearly defined F-actin network in *Chlamydomonas* is the fertilization tubule. The fertilization tubule is an F-actin-rich structure found in mating type plus gametes (Detmers et al., 1985; Detmers et al., 1983), which during mating protrudes from the ‘doublet zone’, a region between the two flagella (Detmers et al., 1983). Phallacidin staining of F-actin strongly labels fertilization tubules (Detmers et al., 1985), and isolation of fertilization tubules has revealed actin as a major component (Wilson et al., 1997). Additionally, null mutants lacking conventional actin cannot form fertilization tubules (Kato-Minoura, 1997). The context-dependent formation of this well-defined F-actin structure in *Chlamydomonas* provides an exceptional opportunity to understand how a cell is capable of precisely regulating its actin cytoskeleton so that actin polymerization occurs only at a very specific place and time.

*Chlamydomonas* expresses a profilin (PRF1) that, like other profilins, inhibits the nucleation of actin monomers, preventing unwanted actin assembly (Kovar et al., 2001). We have identified a *Chlamydomonas* formin (FOR1) actin assembly factor, which has not been characterized and its cellular role in *Chlamydomonas* not yet determined. Therefore, we sought to characterize the formin FOR1 and determine how FOR1 assembles actin monomers bound to PRF1. Additionally, we wished to determine the role of FOR1 in *Chlamydomonas* cells. We found that in addition to inhibiting nucleation, PRF1 potently inhibits the barbed end elongation of actin filaments at relatively low concentrations. However, FOR1 overcomes this inhibition and swiftly assembles PRF1-bound actin monomers into actin filaments that elongate rapidly. *Chlamydomonas* cells treated with the formin inhibitor SMIFH2 do not form fertilization tubules, nor do *for1* or *prf1-1* mutants, suggesting that the collective activities of PRF1 and FOR1 regulate acute F-actin assembly for mating in *Chlamydomonas*.

## RESULTS

### PRF1 inhibits nucleation and elongation of actin filaments

Plant, fungal, and metazoan cells maintain a large pool of unassembled G-actin bound to profilin (Carlsson et al., 1977; Kaiser et al., 1999; Lu and Pollard, 2001). Profilin inhibits the unwanted nucleation of new actin filaments, but once an actin filament has been formed, profilin-bound actin monomers add to the barbed end of growing actin filaments to essentially the same degree as free monomers (Pollard and Cooper, 1984). Additionally, mammalian profilins promote nucleotide exchange (such as ADP to ATP) of actin, although plant profilins do not (Goldschmidt-Clermont et al., 1991; Mockrin and Korn, 1980; Perelroizen et al., 1996; Perelroizen et al., 1994).

*Chlamydomonas* profilin PRF1 is found throughout the cytoplasm and flagellar compartments of the cell, but is enriched at the base of the flagella in vegetative cells and below the fertilization tubule in mating type plus gametes (Kovar et al., 2001). Unlike typical profilins, *Chlamydomonas* PRF1 inhibits the nucleotide exchange of bound G-actin (Kovar et al., 2001). PRF1 might therefore inhibit actin assembly in cells more potently than other profilins. We confirmed that the spontaneous assembly of actin monomers was inhibited by PRF1 in a concentration-dependent manner (Figure 1A), like other profilins including fission yeast SpPRF, revealing a relatively high affinity for actin monomers (*K*_d_=0.14 μM). Surprisingly, by directly observing the spontaneous assembly of 1.5 μM Mg-ATP actin monomers using Total Internal Reflection Fluorescence (TIRF) microscopy, we found that unlike other profilins such as SpPRF, PRF1 also significantly inhibits the barbed end elongation of actin filaments at concentrations where the ratio of PRF1 to actin is equal or only 2- to 3-fold higher (Figure 1B). Therefore, PRF1 is a multi-faceted inhibitor of actin polymerization that potently prevents both actin filament nucleation and elongation.

**Figure 1:**
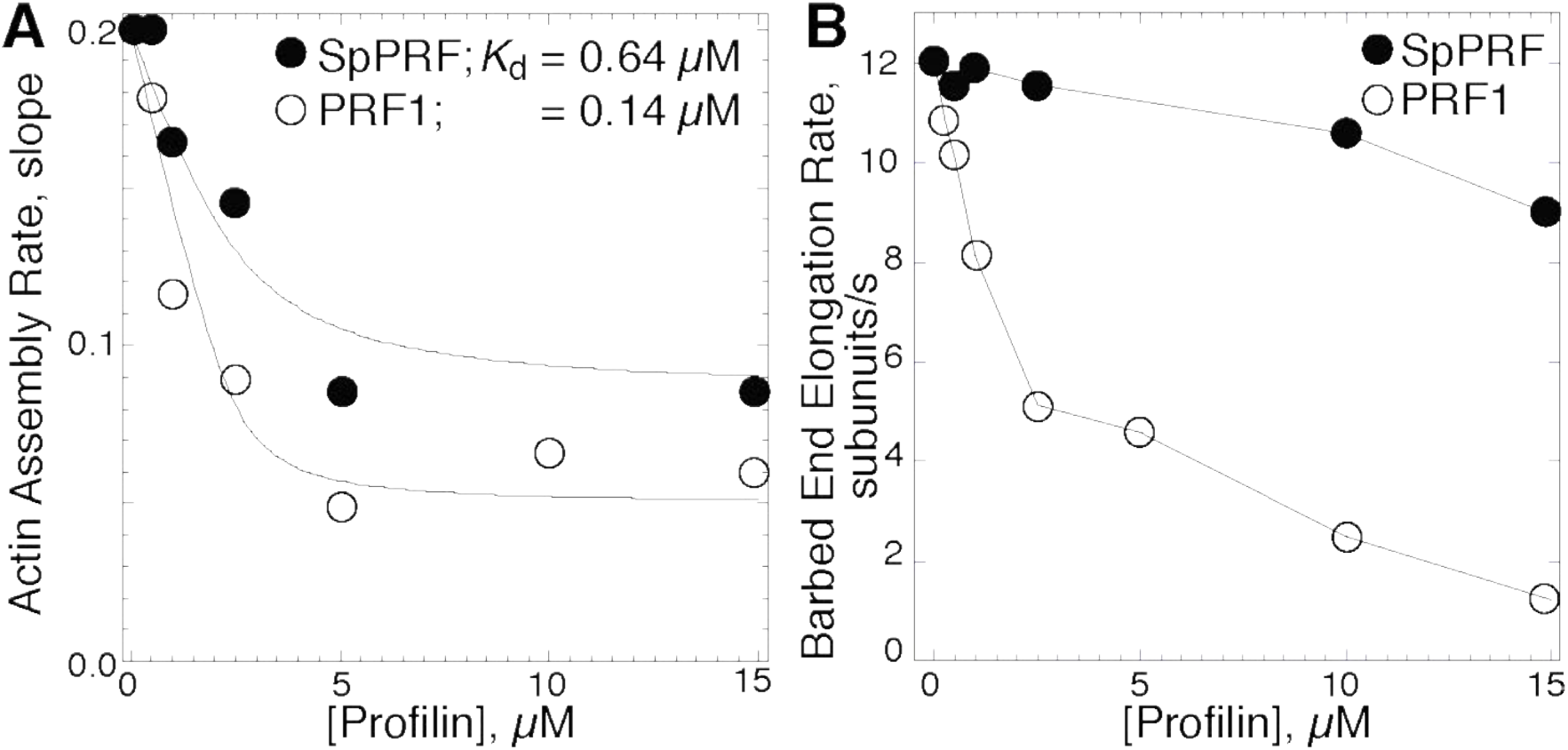
PRF1 inhibits nucleation and elongation of actin filaments. (A) Slopes of spontaneous pyrene actin assembly assays (1.5 *μ*M Mg-ATP actin, 20% pyrene labeled) with increasing concentrations of fission yeast profilin SpPRF or *Chlamydomonas reinhardtii* profilin PRF1. Curve fits reveal affinities of SpPRF and PRF1 for actin monomer. (B) Barbed end elongation rates of 1.5 *μ*M Mg-ATP actin (10% Alexa-488 labeled) in the presence of increasing concentrations of SpPRF or PRF1, measured by TIRF microscopy.

An inhibitory profilin such as PRF1 is ideal to prevent unwanted spontaneous actin assembly. However, as F-actin is present within the fertilization tubule during mating, F-actin polymerization must be facilitated at the correct time and place. Therefore, we speculated that an actin assembly factor such as a formin could be responsible for rapid actin assembly at fertilization tubule sites.

### Formin identification in *Chlamydomonas reinhardtii*

A BLAST search for conserved formin FH2 domain lasso and post sequences using mouse formin (CAA37668 – aa 986-1251) as query identified a *Chlamydomonas* gene locus (Cre03.g166700 in the version 5.6 genome assembly) as a candidate formin. Manual inspection of the genome region upstream of the lasso element revealed an FH1 domain containing at least three proline rich repeats (PRRs) in the same reading frame with typical 6-8aa spacing between. An additional 7 PRRs with typical short (8-12aa) spacing were found further upstream of an unusually long spacer of 37 amino acids. A Kazusa DNA Research Institute EST sequence from *Chlamydomonas* (HCL081g04) confirmed splicing of the putative FH2 domain to the first three PRRs of the FH1 domain. A full length cDNA sequence provided by Susan Dutcher (personal communication) confirmed expression of the long spacer and all 10 PRR regions within a 3157 aa protein (Figure 2A). This formin was named *Chlamydomonas reinhardtii* <u>for</u>min <u>1</u> (FOR1). We created bacterial expression constructs containing either 3 or 10 PRRs along with the FH2 domain (FOR1(3P,FH2) or FOR1(10P,FH2), respectively) and confirmed their ability to stimulate actin polymerization (Figure 2), suggesting that the expressed protein is a formin. There are three other FH2 domain containing genes in *Chlamydomonas*, which we have denoted FOR2 (Cre05.g232900), FOR3 (Cre06.g311250), and FOR4 (Cre04.g229163).

**Figure 2:**
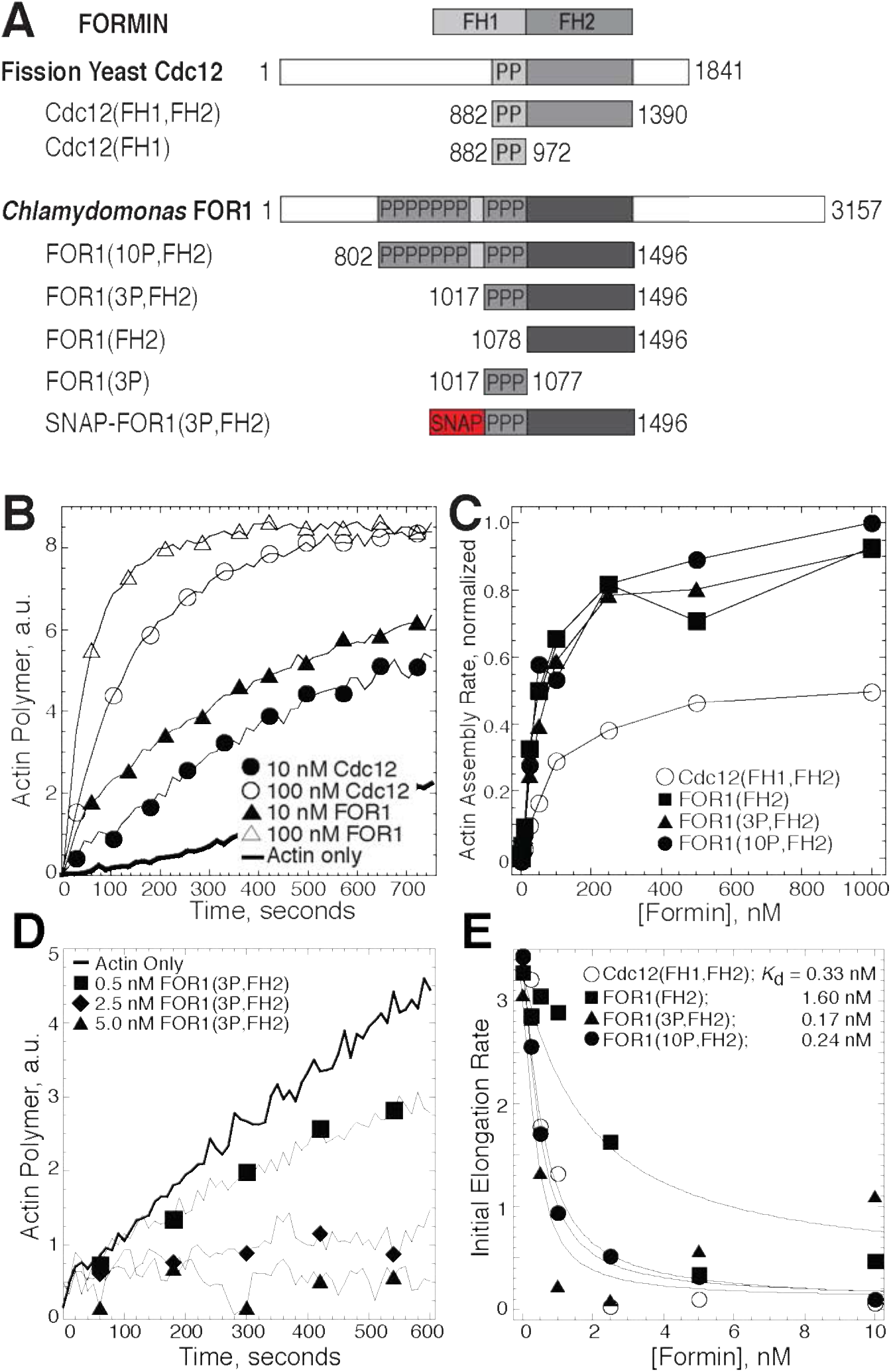
FOR1 efficiently nucleates actin filaments that elongate slowly. (A) Domain organizations and constructs used in this study of fission yeast formin Cdc12 and *Chlamydomonas reinhardtii* formin FOR1. Numbers denote amino acid residues. Each “P” indicates a putative profilin binding site of at least 6 prolines within 8 residues. (B and C) Spontaneous assembly of 2.5 *μ*M Mg-ATP actin monomers (20% pyrene labeled). (B) Pyrene fluorescence over time for actin alone (thick curve), and with 10 (●) or 100 nM (○) Cdc12(FH1,FH2) or 10 (▲) and 100 nM (△) FOR1(3P,FH2). (C) Dependence of the normalized actin assembly rate (slope) on the concentration of Cdc12(FH1,FH2) (○), FOR1(FH2) (■), FOR1(3P,FH2) (▲), and FOR1(10P,FH2) (●). (D and E) Seeded assembly of 0.2 *μ*M Mg-ATP actin monomers (20% pyrene labeled) onto 0.5 μM preassembled filaments. (D) Pyrene fluorescence over time for actin alone (thick line) or in the presence of 0.5 (◼), 1.0 (◆), or 2.5 nM (▲) FOR1(3P,FH2). (E) Dependence of the initial barbed end assembly rate on formin concentration. Curve fits revealed equilibrium dissociation constants of 0.33 nM for Cdc12(FH1,FH2) (○), 1.6 nM for FOR1(FH2) (■), 0.17 nM for FOR1(3P,FH2) (▲), and 0.24 nM for FOR1(10P,FH2) (●).

### FOR1 efficiently nucleates but weakly elongates actin filaments

Formins are a conserved family of actin assembly factors that nucleate actin filaments. Additionally, formins increase the F-actin elongation rate in the presence of profilin by remaining processively associated with the barbed end (Breitsprecher and Goode, 2013). Formins contain actin assembly FH1 and FH2 domains, which are typically flanked by regulatory regions. Functional formins are dimers, with two FH2 domains interacting head-to-tail to create a donut-shaped dimer capable of creating a stable actin ‘nucleus’ (Otomo et al., 2005). In addition, the FH2 dimer remains processively associated with the elongating barbed end of an actin filament (Kovar, 2006). The unstructured FH1 domains are rich in PRRs that bind to profilin and promote rapid association of profilin-actin with the barbed end of an elongating filament. To investigate the actin assembly properties of the formin FOR1, we created a set of constructs containing the FOR1 FH1 and FH2 domains, alone or in combination (Figure 2A). Because our initial inspection of the FOR1 gene suggested that depending upon splicing it contained either 3 or 10 PPRs within its FH1 domain, we created constructs containing either 3 or 10 PPRs. We focused on constructs containing solely the FH1 and FH2 domains as full-length formins are difficult to purify, and active formin constructs containing only the FH1 and FH2 domains have been extensively studied for numerous formins (Breitsprecher and Goode, 2013). However, as the C-terminal domains of certain formins have been shown to affect their actin assembly properties (Gould et al. 2011), it will be interesting in the future to investigate how FOR1 domains modify FOR1 activity.

FOR1’s capacity to stimulate actin assembly in the absence of profilin was initially investigated by measuring the effect of FOR1 on actin polymerization over time using spontaneous pyrene actin assembly assays. FOR1 containing the FH2 domain alone (FOR1(FH2)) or both the FH1 and FH2 domains (FOR1(3P,FH2) or FOR1(10P,FH2)) stimulate actin assembly in a nearly identical, concentration-dependent manner (Figure 2B-C), and more potently than a well-characterized control formin fission yeast Cdc12(FH1,FH2) (Figure 2B-C) (Kovar et al., 2003; Scott et al., 2011). Though these results reveal that FOR1 increases the overall rate of actin polymerization, spontaneous pyrene actin assembly assays are unable to differentiate between an increase in the nucleation and/or elongation of actin filaments.

To differentiate between the contributions of nucleation and elongation to the overall enhanced polymerization rate, we initially examined the effect of FOR1 on actin filament elongation using seeded pyrene actin assembly assays. In the presence of actin filament seeds, elongation of the seeds dominates the reaction and the contribution of nucleation to the overall actin polymerization rate is eliminated. Addition of FOR1(FH2), FOR1(3P,FH2) or FOR1(10P,FH2) to seeded assembly reactions each reduced the actin assembly rate in a concentration dependent matter (Figure 2D-E). This result suggests that FOR1 inhibits actin filament elongation, and that the increased actin assembly rate observed in spontaneous pyrene actin assays is due to FOR1-mediated nucleation. Fits of the initial seeded polymerization rates over a range of formin concentrations revealed dissociation rate constants (*K*_d_) for actin filament barbed ends in the low nanomolar range: FOR1(FH2) (*K*_d_=1.6 nM), FOR1(3P,FH2) (*K*_d_=0.17 nM), FOR1(10P,FH2) (*K*_d_=0.24 nM), and Cdc12(FH1,FH2) (*K*_d_=0.33 nM) (Figure 2E).

### Fission yeast profilin SpPRF enhances FOR1-mediated actin assembly

In the absence of profilin, *Chlamydomonas* formin FOR1 has potent nucleation activity but also significantly inhibits actin filament barbed end elongation, similar to the fission yeast formin Cdc12. However, like other formins (Kovar et al., 2006), Cdc12-associated filaments elongate their barbed ends ~30-fold faster when fission yeast profilin SpPRF is included in the reaction (Kovar et al., 2003; Scott et al., 2011). We hypothesized that profilin would also increase the elongation rate of filaments nucleated by FOR1. We first tested the ability of profilins PRF1 and SpPRF to bind to the FH1 domains of FOR1 and Cdc12. Interestingly, although PRF1 binds much more weakly than SpPRF to non-physiological poly-L-proline (Figure 3A, C) (Kovar et al., 2001), PRF1 and SpPRF have similar affinities for the FH1 domains of both FOR1 and Cdc12 (Figure 3B,C), all with dissociation rate constants (*K*_d_) within the low micromolar range.

**Figure 3:**
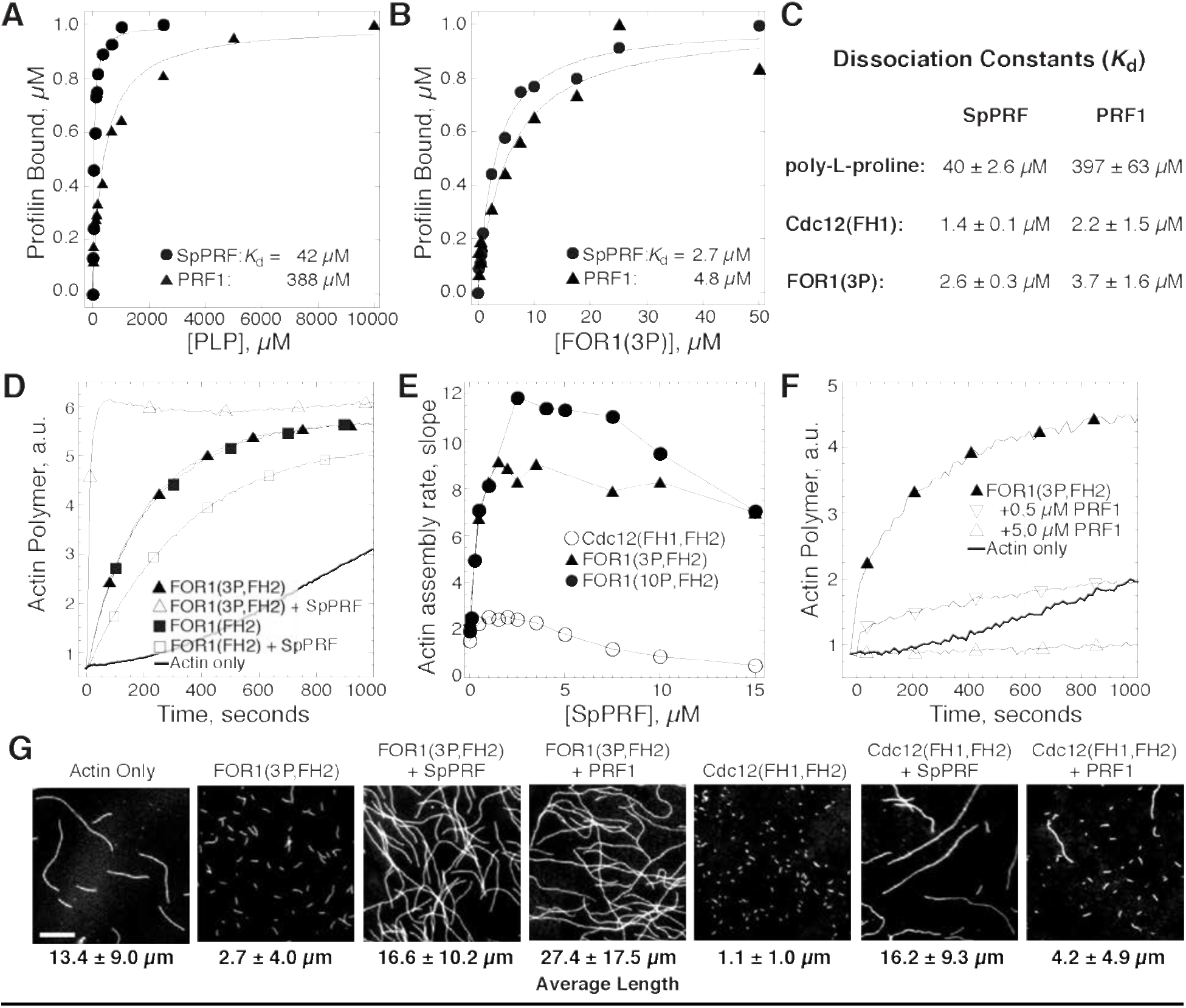
FOR1 stimulates the assembly of profilin-actin. (A-C) Affinity of profilin for poly-L-proline and formin FH1 domains. Dependence of fission yeast SpPRF (⦁) and PRF1 (▲) intrinsic tryptophan fluorescence on the concentration of poly-L-proline (A) and FOR1(3P) (B). (C) Average affinity of SpPRF and PRF1 for poly-L-proline, Cdc12(FH1) and FOR1(3P); n≥3 experiments. (D-F) Spontaneous assembly of 2.5 μM Mg-ATP actin (20% pyrene-labeled). (D) Pyrene fluorescence over time for actin alone (thick curve), and with 10 nM FOR1(FH2) in the absence (◼) or presence (☐) of 2.5 μM SpPRF, and with 10 nM FOR1(3P,FH2) in the absence (▲) or presence (△) of 2.5 μM SpPRF. (E) Dependence of the actin assembly rate (slope) on the concentration of SpPRF for reactions containing 10 nM Cdc12(FH1,FH2) (○), 10 nM FOR1(3P,FH2) (▲) or 10 nM FOR1(10P,FH2) (⦁). (F) Pyrene fluorescence over time for actin alone (thick curve), and with 10 nM FOR1(3P,FH2) in the absence (▲), or presence of 0.5 μM (▿), or 5.0 μM (△) PRF1. (G) Fluorescence micrographs of actin filaments taken 10 minutes after the initiation of the indicated reactions with 10 nM formin and 2.5 μM profilin. Samples were labeled with rhodamine-phalloidin and adsorbed to glass coverslips coated with poly-L-lysine. Scale bar, 5 μm.

Although PRF1 binds well to the FOR1 FH1 domain, the capacity of a formin to add profilin-actin to filament barbed ends depends on complementary interactions with profilin and both the FH1 and FH2 domains of the formin (Bestul et al., 2015; Neidt et al., 2009). We initially tested the ability of FOR1 to assemble SpPRF-actin, as SpPRF is widely compatible with different formin isoforms (Bestul et al., 2015; Neidt et al., 2009). Spontaneous pyrene actin assembly assays revealed that FOR1 constructs containing both the FH1 and FH2 domains (FOR1(3P,FH2) and FOR1(10P,FH2)) rapidly accelerate actin assembly in the presence of SpPRF (Figure 3D-E). Conversely, SpPRF inhibits actin assembly by FOR1(FH2), the construct lacking the FH1 domain (Figure 3D-E). The pyrene actin assembly rates measured for FOR1(3P,FH2) and FOR1(10P,FH2) are significantly greater than those of Cdc12(FH1,FH2) over a range of SpPRF concentrations (Figure 3E), suggesting that SpPRF dramatically increases the processive barbed end elongation rate of FOR1-nucleated actin filaments.

### PRF1-actin is utilized specifically by FOR1

We next examined the ability of FOR1 to assemble actin monomers bound to PRF1. In spontaneous pyrene actin assembly assays, the pyrene fluorescence measured in reactions containing FOR1 and PRF1 is sharply reduced relative to actin alone or actin in the presence of FOR1 (Figure 3F). While this could indicate that PRF1 severely inhibits FOR1-mediated actin assembly, it is also possible that the combination of FOR1 and PRF1 prevents assembly of actin labeled on Cys-374 with pyrene, as we have described for other formin and profilin combinations (Kovar et al., 2006; Scott et al., 2011). Therefore, we directly visualized actin filaments formed in spontaneous pyrene actin assembly assays in the presence of different combinations of formin and profilin. After assembling for 600 seconds, the bulk polymerization reactions were stopped by diluting into TRITC-Phalloidin to allow visualization of filaments by fluorescence microscopy (Figure 3G). In the absence of profilin, FOR1 produces many small actin filaments (average length, 2.7 ± 4.0 μm), indicative of efficient nucleation by FOR1, as suggested by spontaneous pyrene actin assembly assays (Figure 2). Additionally, FOR1 facilitates formation of long actin filaments in the presence of both SpPRF (16.6 ± 10.2 μm) and PRF1 (27.4 ± 17.5 μm). Interestingly, although FOR1 can utilize either SpPRF or PRF1 to elongate actin filaments, Cdc12 is unable to form long filaments in the presence of PRF1 (average length, 4.2 ± 4.9 μm) (Figure 3G), suggesting that PRF1 is tailored for elongation by FOR1. Together, these results indicate that FOR1 is capable of efficient actin filament nucleation, and in the presence of its complementary profilin PRF1, rapidly elongates these filaments. In addition, the inability of Cdc12 to elongate PRF1-associated actin suggests that FOR1 and PRF1 are tailored to precisely and rapidly polymerize F-actin.

### FOR1 rapidly and processively elongates actin filaments in the presence of PRF1

To directly examine the effect of PRF1 on FOR1-mediated actin assembly, we visualized the assembly of 1 μM Mg-ATP actin (10% Alexa-488 labeled) over time using TIRF microscopy. As we found that FOR1(3P,FH2) assembled F-actin to a similar degree as FOR1(10P,FH2) in several actin assembly assays (Figure 2, 3), we chose to use FOR1(3P,FH2) in the remainder of our biochemistry experiments. Actin filaments alone (control) elongate at a rate of 11.5 subunits per second (Figure 4A). In the presence of 1 nM FOR1(3P,FH2), two populations of filaments are observed: actin filaments elongating at the control rate (9.1 sub/s, red arrowheads), and actin filaments elongating at a significantly slower rate (0.3 sub/s, blue arrowheads) (Figure 4B). We predict that the slow-growing filaments are bound at their barbed end by FOR1, which significantly reduces their elongation, while filaments elongating at the control rate are not bound by FOR1 (Kovar et al., 2003, Kovar et al., 2006). In the presence of 1 nM FOR1 and 2.5 μM PRF1, two distinct populations of filaments are again observed: actin filaments elongating at a rate slower than the control rate (4.2 sub/s) and rapidly elongating actin filaments (63.2 sub/s) (Figure 4C). The assembly rate of internal control filaments is slower in these reactions because PRF1 inhibits actin filament elongation (Figure 1B), while FOR1 can efficiently utilize PRF1-bound actin to rapidly elongate actin filaments. The 200-fold difference between the elongation rate of FOR1-bound F-actin in the absence (~0.3 sub/s) and presence of PRF1 (~60 sub/s) is one of the largest observed (Kovar, 2006).

**Figure 4:**
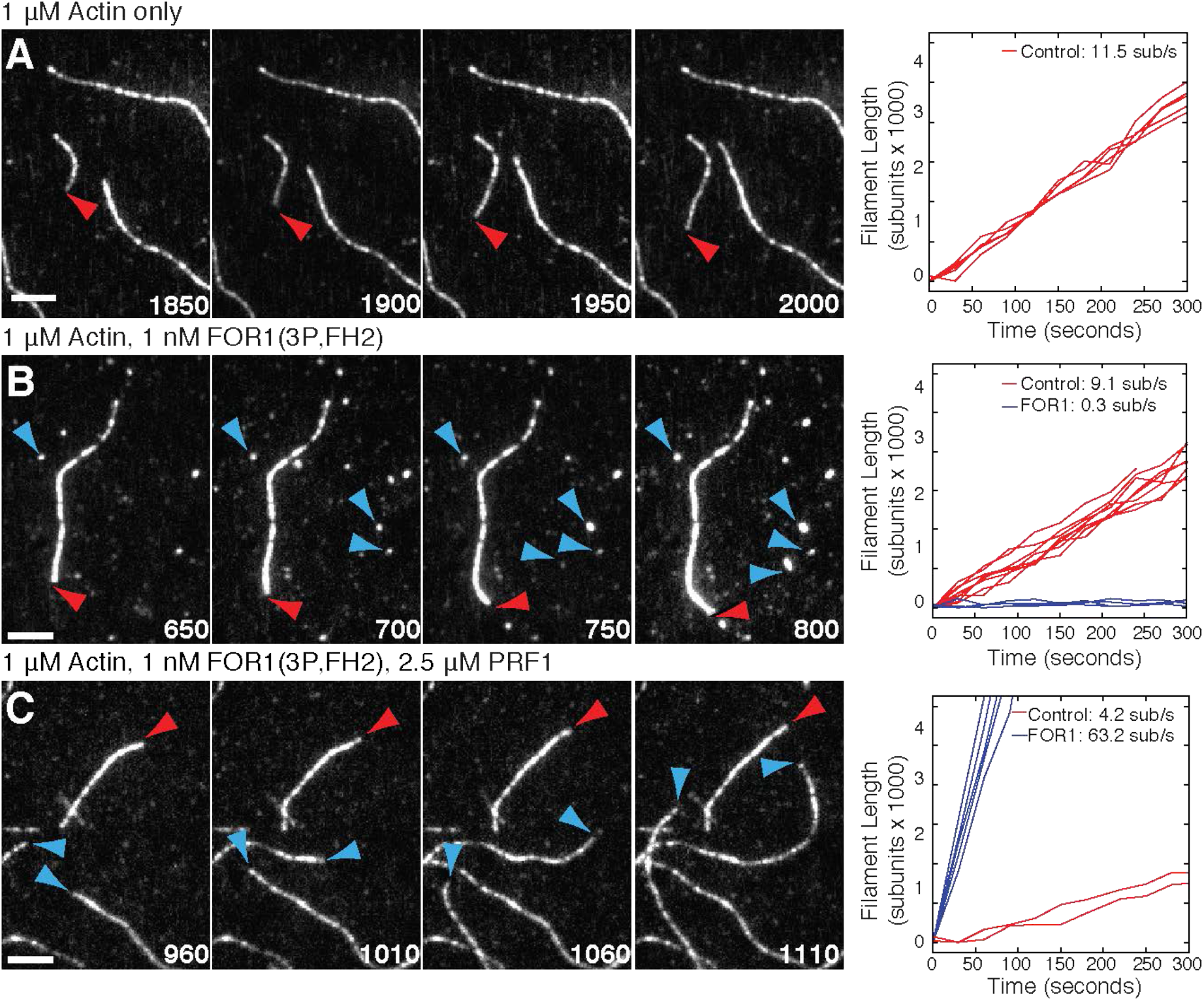
FOR1 rapidly elongates actin filaments in the presence of PRF1. **(A-C)** TIRF microscopy of 1 μM Mg-ATP actin (20% Alexa-488 labeled). (Left) Time lapse micrographs with time in seconds of actin alone **(A)**, with 1 nM FOR1(3P,FH2) **(B)**, or with 1 nM FOR1(3P,FH2) and 2.5 μM PRF1. Red and blue arrowheads denote control (formin independent) and FOR1-dependent filaments, respectively. Scale bars, 5 μm. (Right) Rates of filament growth for control (red lines) and FOR1-associated (blue lines) filaments.

Our results suggest that FOR1 and PRF1 cooperate to rapidly elongate F-actin. To directly visualize and confirm this finding, we made a SNAP-tagged construct of FOR1(3P,FH2) that was labeled with SNAP-549 dye for multi-color TIRF microscopy experiments (Figure 5). In the absence of PRF1, SNAP-FOR1(3P,FH2) remains continuously associated with the barbed end of small, slow growing actin filaments (Figure 5A, blue arrowheads), consistent with our finding that FOR1 can nucleate actin filaments but significantly slows actin filament elongation. Conversely, in the presence of PRF1, SNAP-FOR1(3P,FH2)-associated actin filaments elongate rapidly (Figure 5B,D) compared to control filaments (Figure 5C).

**Figure 5:**
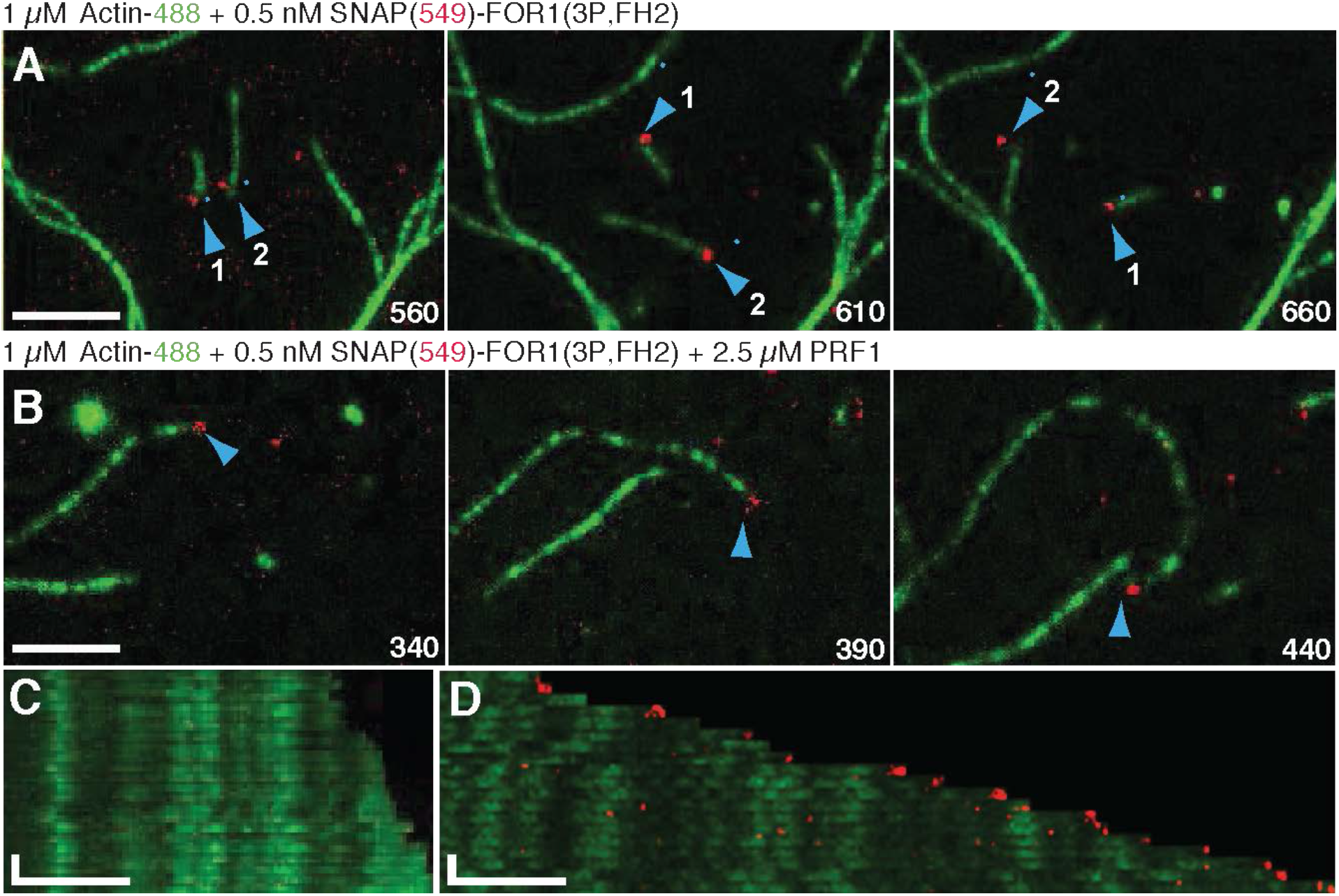
FOR1 is processive on F-actin barbed ends in the absence and presence of PRF1. **(A-D)** Two-color TIRF microscopy of 1 μM Mg-ATP actin (10% Alexa-488 labeled) with 0.5 μM SNAP-FOR1(3P,FH2) (549-labeled) in the presence or absence of 2.5 μM PRF1. Blue arrowheads denote formin-bound filaments. **(A)** 0.5 μM SNAP-FOR1(3P,FH2) alone. **(B)** 0.5 μM SNAP-FOR1(3P,FH2) in the presence of 2.5 μM PRF1. **(C and D)** Kymographs of control **(C)** and formin-bound **(D)** filaments from **(B)**. Scale bars, x-axis, 5 μm. Time bars, y-axis, 30 sec.

### PRF1 favors formin-over Arp2/3 complex-mediated assembly

We found that PRF1 potently prevents spontaneous actin assembly by inhibiting both nucleation and barbed end elongation, whereas FOR1 overcomes this inhibition and facilitates the assembly of rapidly elongating actin filaments. In addition to enhancing formin-mediated elongation, profilin has also been shown to tune F-actin network formation by inhibiting Arp2/3 complex-mediated actin filament branch formation (Rotty et al., 2015; Suarez et al., 2015). We speculated that PRF1 might be a particularly potent inhibitor of Arp2/3 complex by inhibiting both branch formation and subsequent elongation. We tested this possibility by performing biomimetic assays in which fission yeast Arp2/3 complex activator Wsp1 (Figure 6A, Movie 1) or formin FOR1 (Figure 6C, Movie 2) are attached to a polystyrene bead within a standard TIRF microscopy chamber. We sequentially flowed actin alone followed by PRF1-bound actin into the microscopy chamber to assess the effect of PRF1 on formin- and Arp2/3 complex-mediated actin assembly. FOR1-bound beads poorly assembled F-actin in the absence of profilin (Figure 6D(1), E(1)), similar to what we observed in standard TIRF microscopy assays (Figure 4B, 5A). Addition of PRF1-actin into the TIRF chamber triggered rapid actin filament assembly (Figure 6D(2), E(2)). Photobleaching the rapidly assembling F-actin showed a recurrence of high fluorescence at the bead (Figure 6D(3), F), indicative of rapid actin filament assembly by FOR1 at the bead surface. Conversely, beads coated with Wsp1 assembled branched actin filaments normally following incubation with actin and Arp2/3 complex (Figure 6A(1), E(1)). However, filament growth was halted following addition of new actin and Arp2/3 complex with PRF1 (Figure 6B(2), E(2)). Photobleaching of F-actin revealed very little new F-actin assembly at barbed ends, consistent with PRF1 inhibition of actin filament elongation. In addition, very little F-actin assembly occurred at the bead, demonstrating inhibition of Arp2/3 complex-mediated branch formation at the bead surface (Figure 6B(3), F).

**Figure 6:**
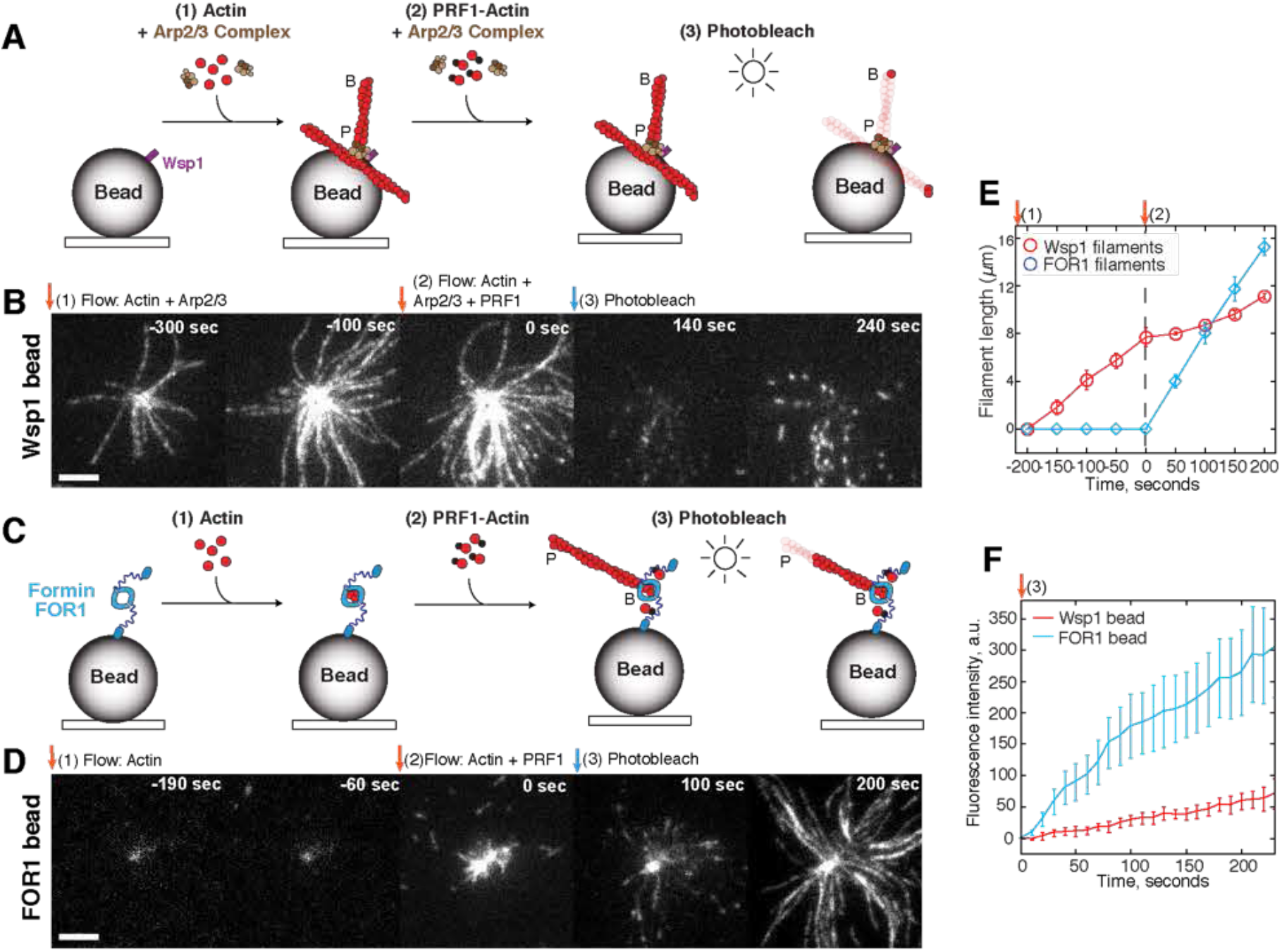
PRF1 facilitates formin-over Arp2/3 complex-mediated assembly. **(A-F)** TIRF microscopy bead assays. Fission yeast Arp2/3 complex activator Wsp1 or formin SNAP-FOR1(3P,FH2) (unlabeled) are adsorbed to a polystyrene bead and the effect of PRF1-actin on network formation is observed. ‘B’ and ‘P’ indicate actin filament barbed and pointed ends, respectively. Scale bars, 5 μm. **(A-B)** Reactions containing beads coated with Wsp1. 1.5 μM Mg-ATP actin (10% Alexa-488 labeled) and 30 nM Arp2/3 complex is initially flowed into the chamber (1), followed by actin, Arp2/3 complex, and 2.5 μM PRF1 (2). Filaments are then photobleached to observe new F-actin assembly (3). **(C-D)** Reactions containing beads coated with FOR1. Actin is initially flowed into the chamber (1), followed by actin and PRF1 (2), and then photobleached (3). **(E)** Actin filament length over time for filaments associated with Wsp1 (red) or FOR1 (blue) beads. The timepoints of initial flow of actin (1) and actin with PRF1 (2) are indicated. Value reported is mean ± s.e.m., n=5 filaments. **(F)** Quantification of fluorescence intensity (actin assembly) at the surface of Wsp1-coated (red) or FOR1-coated (blue) beads following flow-in of PRF1-actin and photobleaching. Each experiment was replicated twice. Value reported is mean ± s.e.m., n=3 Wsp1 or n=4 FOR1 beads.

### Fertilization tubule formation is prevented by the formin inhibitor SMIFH2

PRF1 is a potent inhibitor of actin filament nucleation and elongation. However, PRF1-bound actin can be rapidly assembled by FOR1. We were interested in the role that this tailored protein interaction plays in facilitating actin polymerization *in vivo*. As the fertilization tubule in *Chlamydomonas* is known to be F-actin rich and appears by EM to contain a parallel array of linear actin filaments (Detmers et al., 1983), we suspected that a formin like FOR1 might assemble the long actin filaments required for fertilization tubule formation in *Chlamydomonas* gametes. To test this, we first chemically induced fertilization tubule formation in gametes and stained cells with fluorescent phalloidin to label F-actin (Figure 7B-H). Fertilization tubules were observed in ~43% of untreated or DMSO (control)-treated induced gametes (Figure 7B, C and H). As expected, treatment with 10 μM latrunculin B, which depolymerizes F-actin networks, eliminated fertilization tubules (Figure 7D, H). We then tested whether chemically inhibiting formins would affect fertilization tubule formation. Formin inhibitor SMIFH2 potently inhibited FOR1-mediated actin assembly *in vitro* (Figure 7A) (Rizvi et al., 2009). Correspondingly, though 10 μM of formin inhibitor SMIFH2 had little effect on tubule formation (Figure 7E, H), only 5% of gametes formed fertilization tubules in the presence of 100 μM SMIFH2 (Figure 7F, H). To confirm that fertilization tubule loss with 100 μM SMIFH2 is specific, we also treated cells with 100 μM of Arp2/3 complex inhibitor CK-666 (Nolen et al., 2009). Similar to controls, ~40% of CK-666 cells formed fertilization tubules (Figure 7G, H), indicating that FOR1-mediated but not Arp2/3 complex-mediated F-actin assembly is required for fertilization tubule formation. Furthermore, we also found that FOR1 is capable of bundling actin filaments to a similar extent as fission yeast formin Fus1, the formin involved in mating projectile formation in fission yeast cells (Figure S1), suggesting that in addition to assembling actin filaments, FOR1 could potentially also be involved in bundling actin filaments in the fertilization tubule.

**Figure 7:**
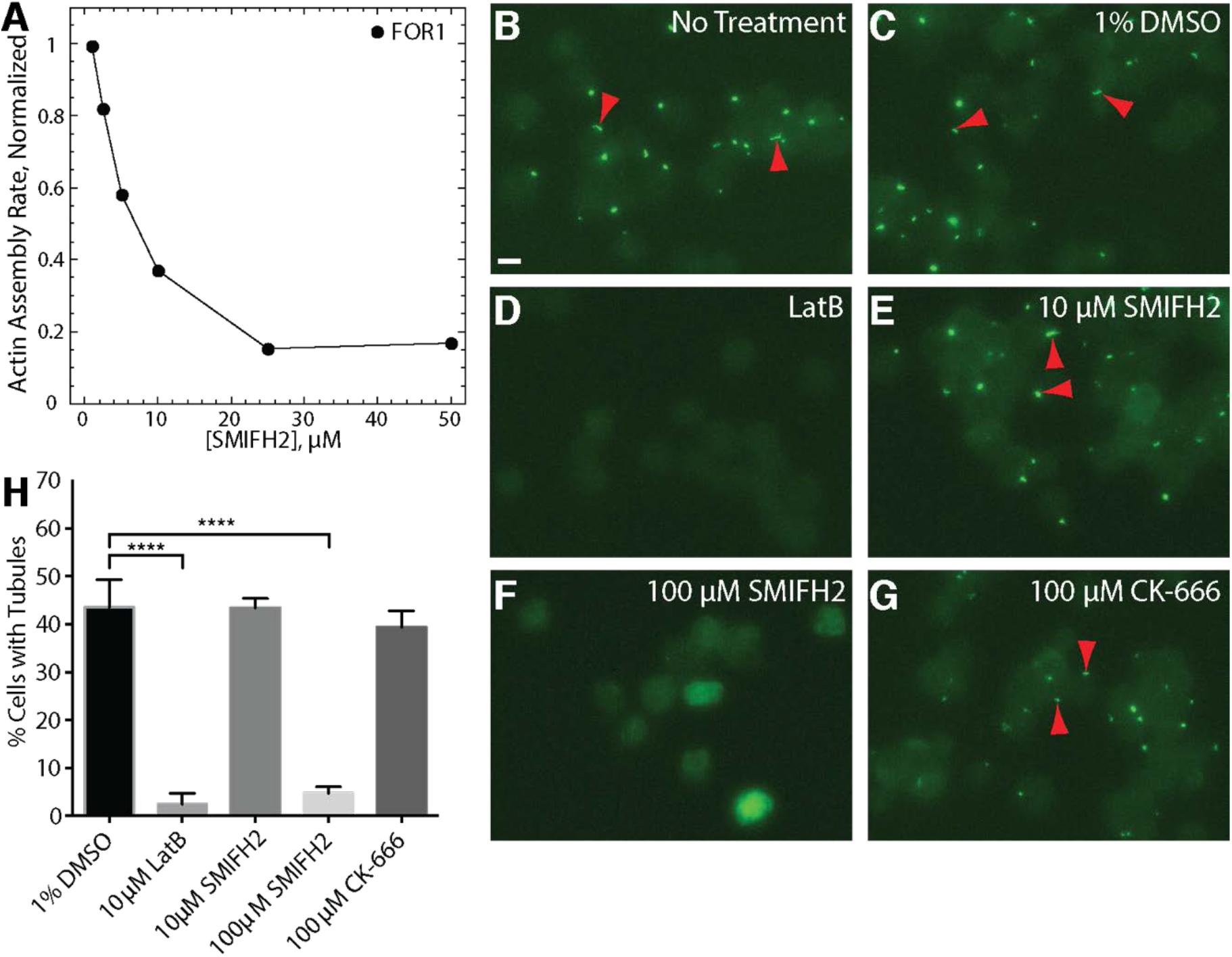
SMIFH2 formin inhibition disrupts fertilization tubules in *Chlamydomonas* gametes. **(A)** Normalized actin assembly rate of FOR1(3P,FH2) (●) in the presence of increasing concentrations of formin inhibitor SMIFH2. **(B-F)** Representative fluorescent micrographs of *Chlamydomonas* gamete fertilization tubules (red arrowheads) labeled with the F-actin marker 488-phalloidin. Scale bar, 10 μm. **(B)** Untreated control. **(C)** 1% DMSO control. **(D)** 10 μM actin depolymerization drug Latrunculin B. **(E)** 10 μM formin inhibitor SMIFH2. **(F)** 100 μM SMIFH2. **(G)** 100 μM Arp2/3 complex inhibitor CK-666. **(H)** Quantification of the percent of cells with fertilization tubules following indicated treatments. n=3 independent experiments. Values reported are mean ± s.d., ****p<0.0001.

### FOR1 and PRF1 mutants fail to form fertilization tubules

SMIFH2 inhibition of fertilization tubule formation suggests that a formin is required to assemble F-actin within the fertilization tubule. However, SMIFH2 is a pan-formin inhibitor that likely has more than one formin target and may have additional non-specific targets as well. As *Chlamydomonas* has four putative formin genes, we wished to determine whether FOR1 specifically assembles F-actin for fertilization tubule formation. Several mutants containing insertions in the FOR1 coding sequence were available from the *Chlamydomonas* mutant library (https://www.chlamylibrary.org). We chose to analyze a mutant with an insertion in exon 3 (Figure 8A, B), upstream of the FH1 and FH2 domains in FOR1. Disruption of FOR1 was confirmed the absence of FH2 domain expression (Figure 8C). In activated wild-type gametes, cells form long fertilization tubules between flagella in 89% of cells (N=200) (Figure 8D). In *for1* insertional mutants, gametes retained their perinuclear actin structures (Figure 8E,E’, and E’’), but began to deform in the region where fertilization tubule extension normally takes place, with no tubule protrusion (Figure 8E’, E’’, white arrows). Sometimes, a filamentous actin focus could be seen at the tip of the deformation (Figure 8E’, white arrowhead). These genetic data confirm that FOR1 is required for normal actin filament assembly in fertilization tubules. In the absence of this formin, attempts to form a tubule cause morphological defects at the apical surface of activated gametes. We also tested the ability of mutants of two other formins to make fertilization tubules. *For2* mutants also exhibited a defect in tubule formation but to a reduced extent than *for1* mutants, as some cells could still make tubules (Figure S2A-E). Because these cells did not show a complete loss of tubules, we tested the functional effects of tubule reduction and found reduced rates of cell fusion (Figure S2F). In contrast, *for3* mutants showed no differences in tubule formation from the wild type (Figure S3). No mutants with an insertion in the coding region of FOR4 were available in the *Chlamydomonas* mutant library. Our data suggest that while FOR2 may also be involved in fertilization tubule formation or maintenance, FOR1 is the primary formin required for proper fertilization tubule formation.

**Figure 8:**
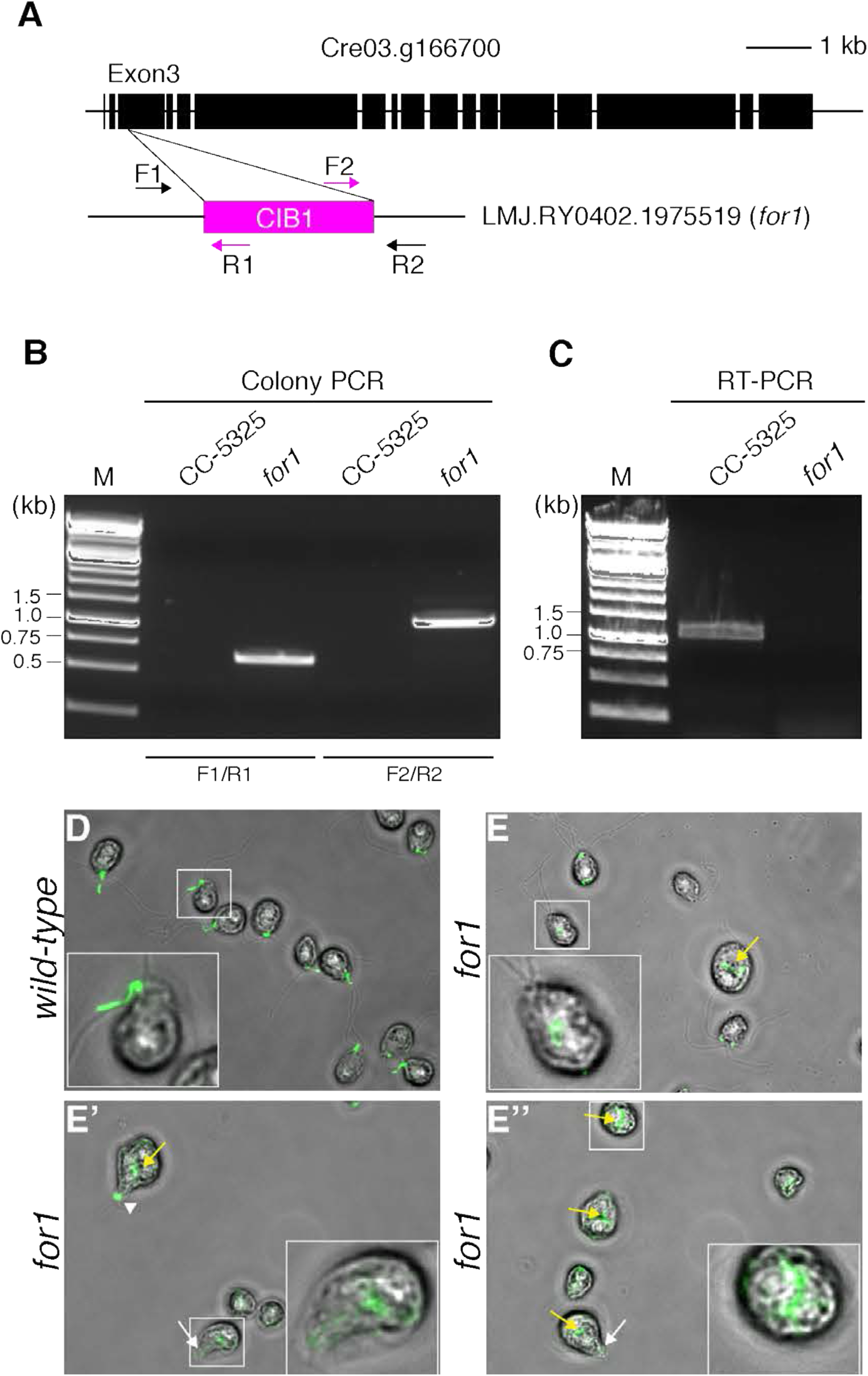
Insertional mutant of *FOR1* fails to make fertilization tubules. **(A)**. Diagram of *Chlamydomonas FOR1* gene. Exons are shown as black rectangles. The CIB1 cassette is inserted in the exon3 of Cre03.g166700 in the *for1* mutant. Arrows indicates primer locations for detecting the cassette insertion. **(B)**. Examination of the genome-cassette junctions by PCR from genomic DNA in the wild type parent strain (CC-5325) or the formin mutant (*for1*). **(C)**. RT-PCR of the functional domain, FH2, of formin in wild type (CC-5325) and the formin mutant (*for1*). **(D-E’’)**. Phalloidin-Alexa Fluor 488 labeled fertilization tubules in wild-type (D) and *for1* mutant **(E, E’, E’’)** cells. *for1* mutants retain mid-cell actin labeling (yellow arrows) and have apical protrusions where tubules should form (white arrows). A collection of labeled actin could sometimes be seen at the tip of the protrusion (white arrowhead).

To further characterize the importance of the formin-profilin interaction on fertilization tubule formation in vivo, we tested whether a temperature-sensitive mutant of PRF1 (*prf1-1*) (Tulin and Cross, 2014; Onishi et al. 2018) was capable of forming fertilization tubules. It has been previously demonstrated that little PRF1 is detectable even at the permissive temperature in *prf1-1* mutants (Onishi et al., 2018). Correspondingly, *prf1-1* mutants showed a complete inability to generate fertilization tubules compared to wild-type controls at both the permissive and restrictive temperatures (Figure 9). However, PRF1 is thought to protect monomeric conventional actin IDA5 from degradation in *Chlamydomonas*, as IDA5 protein is lost and NAP1 upregulated at the permissive temperature in *prf1-1* mutants (Onishi et al., 2018). Given that IDA5 is required for fertilization tubule formation (Kato-Minoura et al., 1997), the *prf1-1* mutant phenotype may also be attributable to the loss of IDA5. We additionally tested whether a temperature-sensitive mutant of the unconventional actin NAP1 (*nap1*) (Onishi et al., 2016) was capable of forming fertilization tubules. We found that mutants of NAP1 (in which IDA5 expression is abundant) showed no defect in fertilization tubule formation (Figure S4). While NAP1 was previously observed within fertilization tubules using NAP1-specific antibodies (Hirono et al., 2003) this population seems inessential for tubule formation.

**Figure 9:**
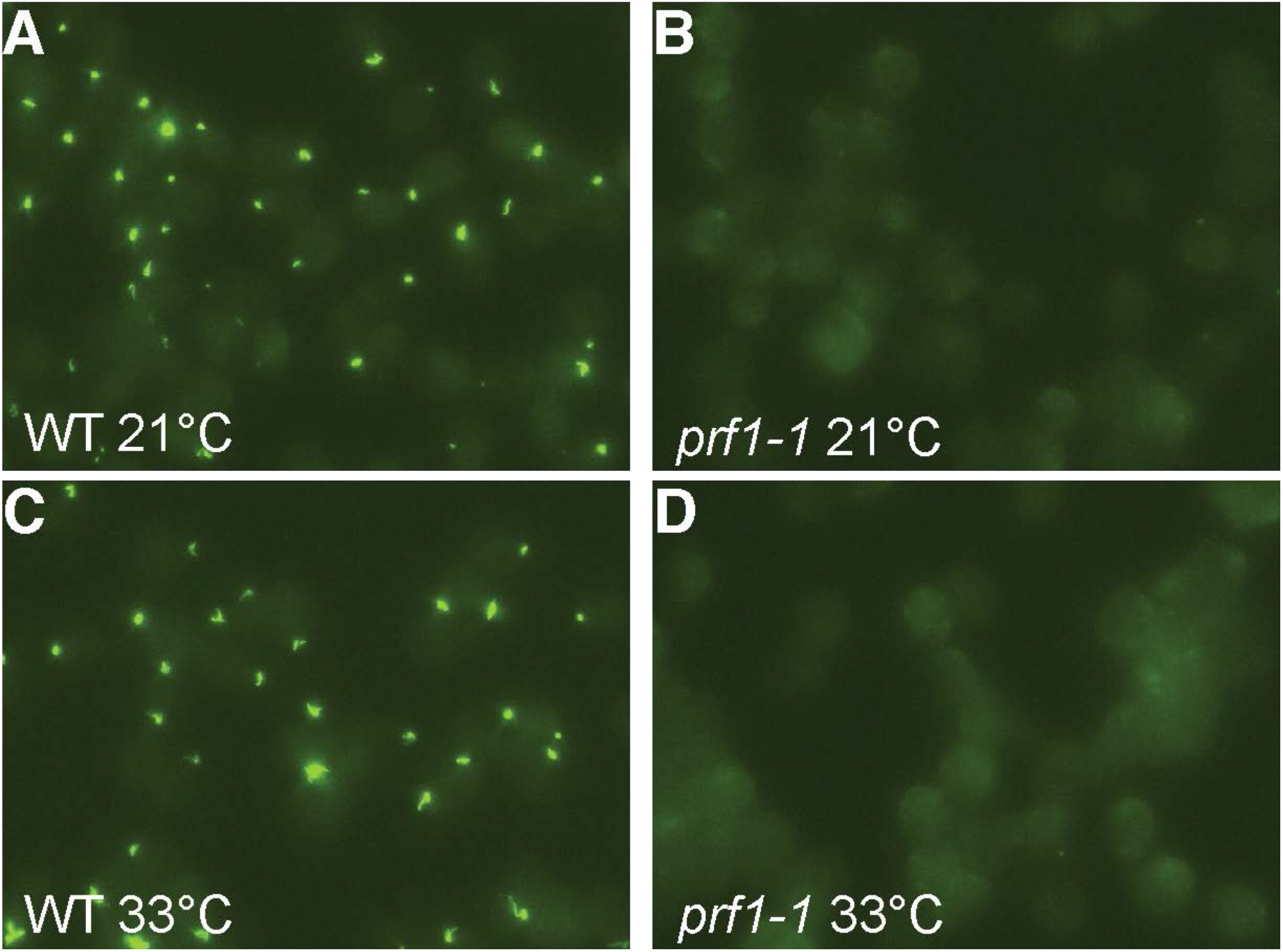
Profilin mutants fail to make fertilization tubules. **(A-D)** Wild-type and profilin mutant (*prf1-1*) cells stained with phalloidin-Alexa Fluor 488 to label filamentous actin-rich fertilization tubules. (A) Wild-type cells at the permissive temperature, 21°C. **(B)** Temperature sensitive *prf1-1* mutants at the permissive temperature, 21°C. **(C)** Wild-type cells at the restrictive temperature, 33°C. **(D)** Temperature sensitive *prf1-1* mutants at the restrictive temperature, 33°C.

## Discussion

### PRF1 as a regulator of F-actin assembly

We found that, unlike typical profilins that primarily inhibit only the nucleation of new actin filaments, PRF1 is an unusual profilin, which, at relatively low concentrations, dramatically prevents actin assembly by inhibiting both the nucleation and barbed end elongation of actin filaments. This effect on actin filament elongation is likely due to an enhanced affinity of PRF1 for the actin filament barbed end. Other profilins have also been shown to decrease barbed end elongation (Pernier et al., 2016, Courtemanche and Pollard, 2013), though at concentrations that are 5-10 times higher than PRF1 (Figure 1B). The enhanced affinity of PRF1 for the barbed end compared to other profilins could make it an ideal tool to study the mechanism and importance of profilin’s competition with other barbed end ABPs (Pernier et al., 2016) such as formin, capping protein, gelsolin, and Ena/VASP. As PRF1 inhibits nucleation, elongation, and the ADP-to-ATP exchange of bound actin monomers (Kovar et al., 2001), PRF1 is a tight regulator of the actin monomer pool, inhibiting spontaneous actin filament assembly in the cell.

Though PRF1 prevents spontaneous actin assembly, FOR1 overcomes the inhibitory effect of PRF1 and utilizes PRF1-bound actin to rapidly assemble actin filaments for the fertilization tubules in mating gametes. We have previously shown that the particular profilin defines the rate of formin-mediated actin assembly (Neidt et al., 2009). The presence of tailored formin-profilin pairs (Bestul et al., 2015) suggests that this interaction is crucial for controlling utilization of an actin monomer pool. The *Chlamydomonas* profilin PRF1 appears to be an extreme example of this, as PRF1-bound actin does not nucleate or elongate well in the absence of FOR1. The other *Chlamydomonas* formins may nucleate and/or elongate PRF1-bound actin to different extents, promoting proper regulation of the profilin-actin pool toward assembly of specific F-actin networks. As our in vitro assays utilize animal actin, it is also possible that the profilin and formins from *Chlamydomonas* interact differently with conventional and/or unconventional *Chlamydomonas* actin than with the animal actin present in our assays. The FH2 domain of formin binds within the hydrophobic cleft between actin subdomains 1 and 3 (Otomo et al. 2005). Of the 11 residues that line the hydrophobic cleft, 100% are conserved in IDA5. In NAP1, 8 of the residues are conserved and 2 are similar amino acids (Figure S5). Of the 21 actin residues that contact profilin (Schutt et al., 1993), 15 are conserved in NAP1 and an additional three are strongly similar amino acid substitutions. The differences between the unconventional NAP1 and the conventional *Chlamydomonas* actin add another layer to the complex regulation of actin assembly in this organism. Future work will involve deciphering how the formins utilize both the conventional and unconventional actins in *Chlamydomonas* to promote specific cellular processes.

Conversely, PRF1 inhibits Arp2/3 complex-mediated actin assembly (Figure 6). Inhibition of Arp2/3 complex-mediated branch formation and elongation could further bias *Chlamydomonas* towards FOR1-mediated assembly, by preventing competition for actin monomers (Suarez and Kovar, 2016; Suarez et al., 2015). *Chlamydomonas* contains ARP2 and ARP3, but its activators have not been identified (Kollmar et al., 2012). The Arp2/3 complex may be involved in assembly and maintenance of the F-actin involved in flagellar membrane or protein trafficking, as treatment with Arp2/3 complex inhibitor CK-666 induces flagellar shortening (Avasthi et al., 2014). The presence of multiple potential F-actin networks provides the possibility that other F-actin assembly factors are also present in *Chlamydomonas*. If so, PRF1 may be involved in regulating competition for actin monomer by these different assembly factors.

In addition to regulating a balance between FOR1- and Arp2/3 complex-mediated F-actin assembly, PRF1 also likely regulates both conventional actin and NAP1 dynamics. Both SMIFH2 (data not shown) and CK-666 (Avasthi et al., 2014) affect flagellar length in mutants lacking conventional actin in which NAP1 is upregulated. This finding suggests that both formin and Arp2/3 complex can nucleate NAP1 filaments. Unlike SMIFH2 and CK-666, latrunculin and cytochalasin do not affect NAP1, which may explain why those treatments do not inhibit cytokinesis. Future work will involve determining the nature of the F-actin networks involved in cytokinesis and flagellar protein trafficking as well as PRF1’s role in ensuring proper F-actin distribution to each network.

### FOR1 in fertilization tubule formation

FOR1 appears to be required for fertilization tubule formation as both *for1* mutants and wild-type gametes treated with the formin inhibitor SMIFH2 do not form fertilization tubules. Fertilization tubule formation in *Chlamydomonas* occurs near the membrane at a site between the two flagella. Prior to fertilization tubule formation, this site is characterized by two parallel electron-dense regions called the membrane zone (immediately adjacent to the membrane) and doublet zone (slightly interior) (Detmers et al., 1983; Goodenough and Weiss, 1975). In a mature fertilization tubule, the pointed ends of actin filaments are attached at the doublet zone (Detmers et al., 1983) while the membrane zone is present at the far end of the extended fertilization tubule, near the F-actin barbed ends. As formins are frequently membrane-anchored, FOR1 is potentially localized to the membrane zone, which extends away from the doublet zone following F-actin formation. FOR1 could additionally be important for bundling the actin filaments in the fertilization tubule (Figure S1), creating a stable projection. Future work will involve determining the factors that regulate FOR1 activity and other ABPs that are involved in proper organization of F-actin at that site.

## ACKNOWLEDGEMENTS

We thank Susan Dutcher (Washington University in St. Louis) and Bill Snell (UT Southwestern) for helpful discussions, including confirmation that *Chlamydomonas* expresses FOR1. We thank Megan Rhyne (University of Tennessee, Knoxville) for helpful comments. This work was supported by National Institutes of Health grants R01 GM079265 (to D.R.K.), P20 GM104936 (to P.A.), NSF Graduate Student Fellowship DGE-1144082 (to J.R.C.), Molecular and Cellular Biology Training Grant T32 GM007183 (to J.R.C. and C.T.S.), and MSTP Training Grant T32GM007281 (to M.J.G.).

## AUTHOR CONTRIBUTIONS

Conceptualization: JRC, MJG, LJM, PA, DRK; Methodology: JRC, MJG, LJM, DRK, PA; Data collection and analysis: JRC, MJG, EWC, DMM, YL, JAS, CTS, SH; Writing-original draft preparation: JRC, MG, PA; Writing-reviewing and editing: JRC, EWC, DMM, PA, DRK; Funding acquisition: JRC, DRK, PA; Resources: LJM, DRK, PA; Supervision: DRK, PA.

## METHODS

### Plasmid construction

Constructs containing different components of the formin actin assembly domains (FH1 and FH2) were prepared for bacterial expression. The preparation of Cdc12(FH1FH2) and Cdc12(FH1) constructs has been described (Neidt et al., 2009). The FOR1 domain constructs were designed based on sequence analysis of the *Chlamydomonas* genome, and Expressed Sequence Tag analysis by Susan Dutcher (Washington University, St. Louis), and were optimized for bacterial expression and custom synthesized (DNA 2.0, Newark, California). Constructs were designed by SnapGene software (from GSL Biotech; available at snapgene.com). All constructs were prepared by standard cloning procedures, consisting of PCR amplification (iProof, Bio-Rad Laboratories) from the commercially prepared DNA. Restriction enzyme cleavage sites and 6x His sequences were included in the reverse primers. PCR products were cloned using restriction enzymes into pET21a (EMD Biosciences) for expression. All amplified sequences were confirmed by sequencing.

### Protein purification

All constructs of FOR1 and PRF1 were expressed in BL21-Codon Plus (DE3)-RP (Agilent Technologies, Santa Clara, CA). Cdc12(FH1FH2) (Kovar and Pollard, 2004), SpPRF (Lu and Pollard, 2001), SpFus1 (Scott et al., 2011), and PRF1 (Kovar et al., 2001) were purified as described previously. FOR1 constructs were His-tag affinity purified. FOR1 constructs were expressed with 0.5 mM isopropyl ß-D-thiogalactopyranoside (IPTG; Sigma-Aldrich) for 16 hours at 16°C. Cells were resuspended in extraction buffer (50 mM NaH_2_PO_4_, pH 8.0, 500 mM NaCl, 10% glycerol, 10 mM imidazole, 10 mM betamercaptoethanol [ßME]) supplemented with 0.5 mM phenylmethylsulfonyl fluoride (PMSF) and protease inhibitors, sonicated, and homogenized in an Emulsiflex-C3 (Avestin, Ottawa, ON, Canada). The homogenate was spun and clarified at 30,000*g* for 15 minutes, then 50,000*g* for 30 minutes and incubated with Talon Metal Affinity Resin (Clontech, Mountain View, CA) for 1 hour at 4°C. The resin was loaded onto a disposable column and washed with 50 mL wash with extraction buffer. FOR1 was then eluted with Talon elution buffer (50 mM NaH_2_PO_4_, pH 8.0, 500 mM NaCl, 10% glycerol, 250 mM imidazole, 10 mM ßME) and dialyzed into formin buffer (20 mM HEPES, pH 7.4, 1 mM EDTA, 200 mM KCl, 0.01% NaN3, and 1 mM DTT).

A_280_ of purified proteins was taken using a Nanodrop 2000c Spectrophotometer (Thermo-Scientific, Waltham, MA). Protein concentrations were determined based on extinction coefficients estimated from amino acid sequences using ProtParam (http://web.expasy.org/protparam/), or from previous studies: PRF1: 19,190 M^−1^ (Kovar et al., 2001), SpPRF: 20,065 M^−1^ (Lu and Pollard, 2001), FOR1(10P,FH2): 29,450 M^−1^, FOR1(3P,FH2): 24,200 M^−1^, FOR1(FH2): 24,400 M^−1^, SNAP-FOR1(3P,FH2): 44,920 M^−1^, and Cdc12 (FH1,FH2): 51,255 M^−1^ (Kovar et al., 2003). Protein concentrations of FH1 constructs Cdc12(FH1), FOR1(FH1), and FOR1(3P) were determined by A_205_ in water ([(A_205_FH1− A_205_buffer)/30]/mol wt). Proteins were flash-frozen in liquid nitrogen and kept at −80°C. SNAP-FOR1(3P,FH2) protein was labeled with SNAP-549 dye (New England Biolabs, Ipswich, MA) as per manufacturer’s instructions prior to each TIRF experiment.

Actin was purified from rabbit or chicken skeletal muscle actin as previously described (Spudich and Watt, 1971). For pyrene assembly assays, actin was labeled with N-(1-Pyrene)Iodoacetamide (Life Technologies, Carlsbad, CA) on Cys-374. As the combination of FOR1 in the presence of PRF1 selected against actin labeled on Cys-374, actin labeled with Alexa Fluor 488 on lysines (ThermoFisher Scientific, Waltham, MA) was used for TIRF microscopy experiments.

### Pyrene assembly and disassembly assays

All pyrene assembly and disassembly assays were carried out in a 96-well plate, and the fluorescence of pyrene-actin (excitation at 364 nm and emission at 407 nm) was measured with a Spectramax Gemini XPS (Molecular Devices) or Safire2 (Tecan) fluorescent plate reader as described (Zimmermann et al., 2016). For spontaneous assembly assays, a 15 μM mixture of 20% pyrene-labeled Mg-ATP-actin monomer with 100X anti-foam 204 (0.005%; Sigma) was placed in the upper well of a 96 well non-binding black plate. Formin and/or profilin, 10X KMEI (500 mM KCl, 10 mM MgCl_2_, 10 mM ethylene glycol tetraacetic acid [EGTA], and 100 mM imidazole, pH 7.0), and Mg-Buffer G (2 mM Tris, pH 8.0, 0.2 mM ATP, 0.1 mM MgCl2 and 0.5 mM DTT) were placed in the lower row of the plate. Reactions were initiated by mixing contents of the lower wells the actin monomers in the upper wells with a twelve-channel pipetman (Eppendorf). For pyrene assembly assays involving SMIFH2, SMIFH2 was added to the lower wells containing FOR1 prior to mixing the upper and lower wells.

For seeded assembly assays, 5.0 μM unlabeled Mg-ATP-actin was preassembled in the upper row of the plate, followed by addition of anti-foam, formin and/or profilin, and Mg-Buffer G. A 2.0 μM mixture of 20% pyrene-labeled actin with Mg-Buffer G was placed in the lower plate row. Mixing actin monomers in lower wells with pre-assembled actin filaments in upper wells initiated reactions.

For depolymerization assays, a 5.0 μM mixture of unlabeled and 50% pyrene-labeled Mg-ATP-actin monomers was preassembled in the upper row of the plate for two hours, followed by addition of anti-foam. Formin, 10X KMEI and Mg-Buffer G were placed in the lower plate row. Reactions were initiated by mixing lower wells with upper wells, diluting the pre-assembled filaments to 0.1 μM.

### Profilin FH1 affinity assays

The affinity of profilin for formin(FH1) was determined by measuring the change in profilin’s intrinsic tryptophan fluorescence by excitation at 295 nm and emission at 323 nm (Perelroizen et al., 1994; Petrella et al., 1996). Profilin (1.0 μM) was incubated with a range of poly-L-proline or formin(FH1) concentrations for 30 min, then profilin fluorescence was read in a Safire2 fluorescence plate reader and plotted versus formin(FH1) concentration. The fluorescence of formin(FH1) alone was subtracted from the fluorescence in the presence of profilin. Dissociation constants (*K*_d_) were determined by fitting a quadratic function to the dependence of the concentration of bound profilin on the concentration of formin(FH1).

### Polymerization and depolymerization rate determination

Actin assembly rates were determined from spontaneous assembly reactions by measuring the slopes of actin assembly following the lag phase to 50% of total actin assembly. Assembly rates from preassembled actin seeds were determined by a linear fit to the first 100 seconds of assembly. Depolymerization rates were determined by a linear fit to the first 100-300 seconds of the reaction.

The affinity of FOR1 for barbed ends was determined as previously described (Kovar *et al.*, 2003). We fit the plot of the dependence of the assembly or disassembly rate on formin concentration using the equation V_i_ = V_if_ + (V_ib_ – V_if_) ((*K*_d_ + [ends] + [formin] – √((*K*_d_ + [ends] + [formin])2 – 4[ends][formin])/2[ends])), where V_i_ is the observed elongation or depolymerization rate, V_if_ is the elongation or depolymerization rate of free barbed ends, V_ib_ is the elongation or depolymerization rate of bound barbed ends, [ends] is the concentration of barbed ends, and [formin] is formin concentration. The nucleation efficiency was calculated by dividing the slope of the spontaneous assembly rate by *k*+ in the absence and presence of profilin and dividing by the formin concentration (Kovar et al., 2006). Depolymerization rates are normalized to the rate of actin assembly alone and expressed as a percent of the standard actin assembly rate.

### Fluorescence micrographs (rhodamine phalloidin)

Unlabeled Mg-ATP-actin was assembled as per standard spontaneous assembly reactions. Actin filaments were then incubated with 1 μM TRITC-Phalloidin (Fluka Biochemika, Switzerland) for 5 minutes. Reactions were terminated by diluting assembled filaments in fluorescence buffer (50 mM KCl, 1 mM MgCl_2_, 100 mM DTT, 20 μg/ml catalase, 100 μg/ml glucose oxidase, 3 mg/ml glucose, 0.5% methylcellulose, and 10 mM imidazole, pH 7.0) and were absorbed to coverslips coated with 0.05 μg/μl poly-L-lysine. Fluorescence microscopy images were collected on an Olympus IX-81 microscope and cooled CCD camera (Orca-ER, Hamamatsu).

### Low-speed sedimentation assays

Sedimentation assays were performed as previously described (Zimmermann et al., 2016). 15 μM Mg-ATP actin monomers were spontaneously assembled for 1 hour in 10 mM imidazole, pH 7.0, 50 mM KCl, 5 mM MgCl_2_, 1 mM EGTA, 0.5 mM DTT, 0.2 mM ATP and 90 μM CaCl_2_ to generate F-actin. Filamentous actin was then incubated with FOR1 or SpFus1 for 20 minutes at 25°C and spun at 10,000g at 25°C. Supernatant and pellets were separated by 15% SDS-PAGE gel electrophoresis and stained with Coomassie Blue for 30 minutes, destained for 16 hours and analyzed by densitometry with ImageJ (Schneider et al., 2012; http://imagej.net).

### TIRF microscopy

Time-lapse TIRF microscopy movies were obtained using a iXon EMCCD camera (Andor Technology, Belfast, UK) fitted to an Olympus IX-71 microscope with through-the-objective TIRF illumination as described (Zimmermann et al., 2016). Mg-ATP-actin (10-20% Alexa-488 labeled) was mixed with a polymerization mix (10 mM imidazole (pH 7.0), 50 mM KCl, 1 mM MgCl2, 1 mM EGTA, 50 mM DTT, 0.2 mM ATP, 50 μM CaCl2, 15 mM glucose, 20 μg/mL catalase, 100 μg/mL glucose oxidase, and 0.5% (400 centipoise) methylcellulose) to induce F-actin assembly (Winkelman et al., 2014). Where stated, formin or profilin was added to the polymerization mix prior to mixing with actin and initiating F-actin polymerization. The mixture was then added to a flow chamber and imaged at 10 s intervals at room temperature. For bead assays, Wsp1 and formin beads were prepared as previously described (Loisel et al., 1999). Carboxylated Polybeads (Polysciences, Warrington, PA) were coated with Wsp1 or unlabeled SNAP-FOR1(3P,FH2) and flowed into the TIRF chamber prior to initiating the reaction.

### Fertilization tubule assay

Wild type 137c (CC-125 mt+) *Chlamydomonas reinhardtii* cells were obtained from the Chlamydomonas Resource Center (University of Minnesota). To induce gametogenesis, cells were grown in M-N (M1 minimal media without nitrogen) overnight under growth lighting. Gametes were mixed with dibutyryl cAMP (13.5mM) and papaverine (135μM) to induce fertilization tubule formation along with different inhibitor preparations; untreated, 1% DMSO (solvent for all inhibitors), 10μM Latrunculin B, 10μM SMIFH2, 100μM SMIFH2, and 100μM CK-666. Cells were placed on a rotator under a LumiBar LED light source (LumiGrow, Inc) for 2hrs. After fertilization tubule induction, cells were adhered to coverslips coated with poly-lysine and fixed with 4% paraformaldehyde in 10mM HEPES. They were permeabilized with −20°C acetone, stained with 100nM Alexa Fluor 488 Phalloidin (Life Technologies) according to manufacturer protocols and mounted on slides with Fluoromount-G (Southern Biotech) for imaging. Slides were imaged with a Nikon Ti-S widefield fluorescence microscope using a Plan Achromat 100x/1.25 NA oil immersion objective lens, a QICam fast 1394 CCD digital camera (QImaging) and NIS Elements software.

All cells in multiple fields of view (~50-100 cells per condition) were counted for presence of fertilization tubules using the ImageJ Cell Counter plugin to determine tubule percentage (#tubules/# total cells) × 100. Means and standard deviations are plotted for experiments done in triplicate. Results were analyzed with one way ANOVA and Dunnett’s multiple comparison post hoc test. For fertilization tubule measurements, line segments were drawn onto projected FITC images and fit with splines using ImageJ. n>45 measurements were collected following a pixel to micron ratio conversion for the optical setup and compared using Kruskal-Wallis and Dunn’s multiple comparison tests.

### Colony PCR, RNA extraction, reverse transcription (RT)-PCR

Colony PCR was performed as previously described (Cao et al., 2009). The genome-cassette junctions were amplified via PCR by using different primers set. The primers set F1/R1 or F2/R2 is used to amplify the left or right insert junction, respectively (F1: <u>ATCAGGAGCCCCCTGTATTT;</u> R1: <u>GCACCAATCATGTCAAGCCT;</u> F2: <u>GACGTTACAGCACACCCTTG;</u> R2: <u>CACCTGACGTGTTGTTGACC</u>). Total RNA was isolated with PureLink™ RNA Mini Kit (Thermo Fisher Scientific). To avoid genomic DNA contamination, on-column PureLink™ DNase (Thermo Fisher Scientific) treatment was performed. First-strand cDNA was synthesized from 1⎧g purified total RNA with SuperScript™ III First-Strand Synthesis system (Thermo Fisher Scientific). For RT-PCR, cDNA fragment coding the majority of the formin FH2 domain was amplified using gene-specific primers For1_F and For1_R (For1_F: <u>CTCCCCCTCCGGTTATGAG;</u> For1_R: <u>CAGACAGCTCGTTCAGCTTG</u>). For both colony PCR and RT-PCR, Phusion® High-Fidelity DNA Polymerase (New England Biolabs) was used. And the amplification conditions were as follows: 98°C for 30 sec, followed by 35 cycles of 98°C for 10 sec, 65°C for 10 sec and 72°C for 60 sec.

