## Supplementary Materials for "*Chlamydomonas reinhardtii* formin FOR1 and profilin PRF1 are optimized for acute rapid actin filament assembly"

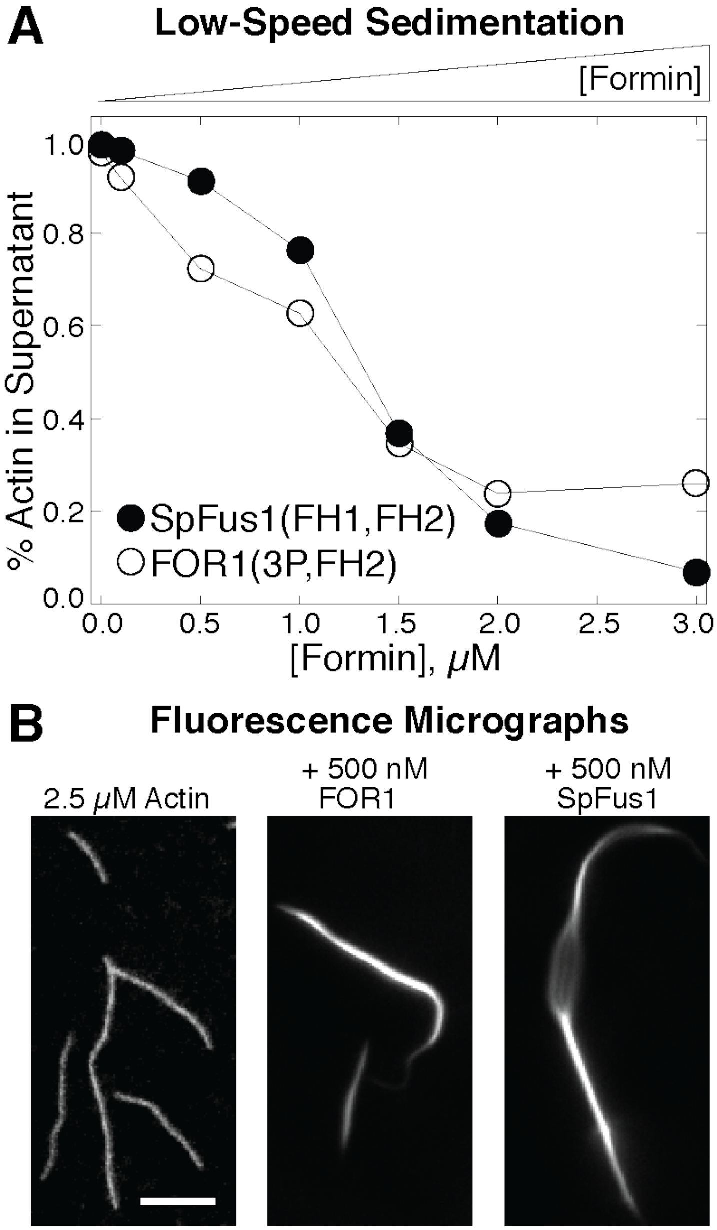


**Supplementary Figure S1: FOR1 bundles F-actin.**

**(A)** Low speed (10,000 x *g*) sedimentation of F-actin preassembled from 5 μM Mg-ATP actin with a range of concentrations of FOR1 (○) or fission yeast formin SpFus1 (●). Plot of the dependence of F-actin in the pellets on the concentration of FOR1 or SpFus1. **(B)** Fluorescence micrographs of F-actin preassembled alone or in the presence of FOR1 or SpFus1 for 20 min and stained with rhodamine-phalloidin. Scale bar, 5 μm.

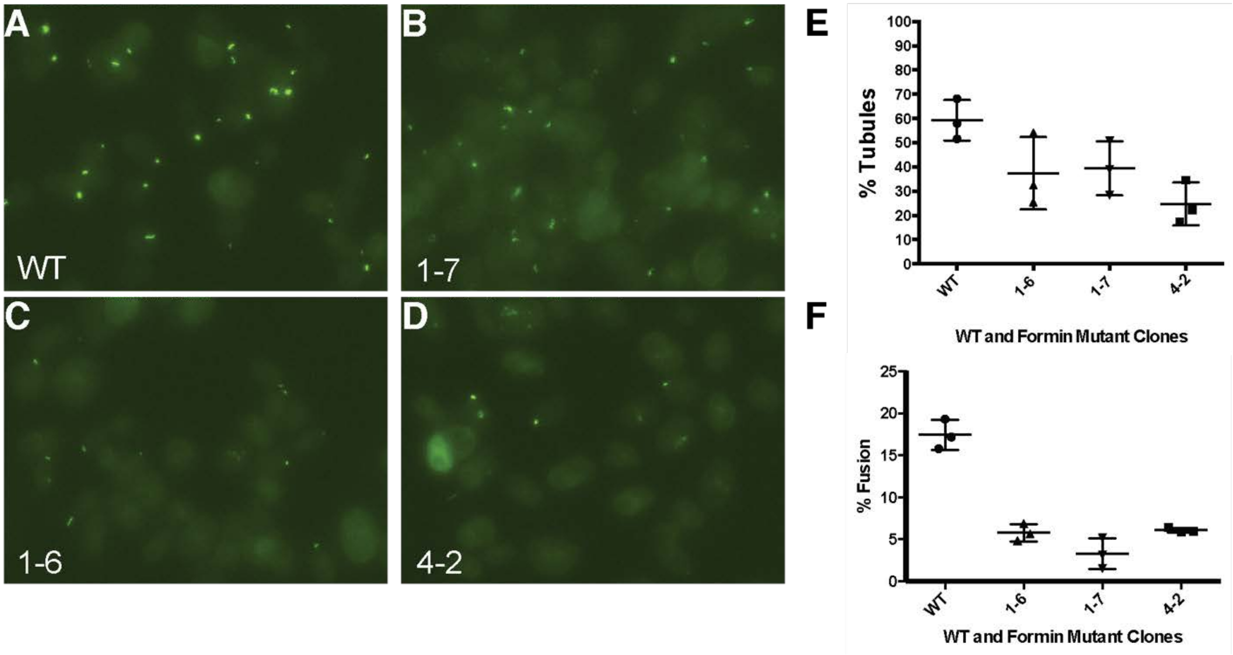


**Supplementary Figure S2: Mutants of FOR2 have reduced rates of fertilization tubule formation and cell fusion.**

**(A-D)** Wild-type cells **(A)** and *for2* mutant isolates 1-7 **(B)**, 1-6 **(C)**, and 4-2 **(D)** stained with phalloidin-Alexa Fluor 488 to label actin-rich fertilization tubules. **(E)** Quantification of percentage of wild-type and *for2* cells that make fertilization tubules. **(F)** Quantification of percentage of wild-type and *for2* cells that fuse during mating.

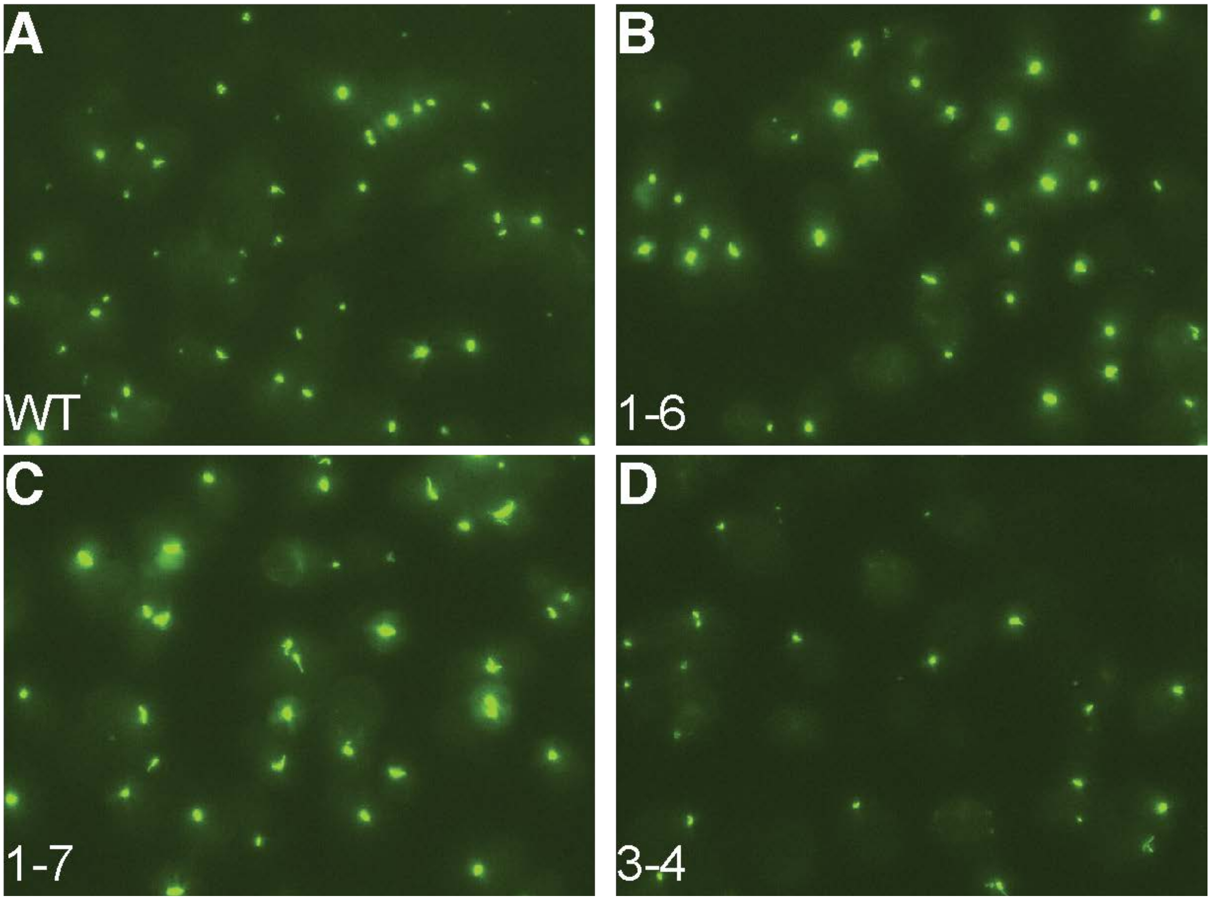


**Supplementary Figure S3: Mutants of FOR3 have no reduction in fertilization tubule formation.**

**(A-D)** Wild-type cells, 74% with fertilization tubules **(A)** and *for3* mutant isolates 1-6 with 77% fertilization tubules **(B)**, 1-7 with 74% fertilization tubules **(C)**, and 3-4 with 72% fertilization tubules **(D)** stained with phalloidin-Alexa Fluor 488 to label actin-rich fertilization tubules.

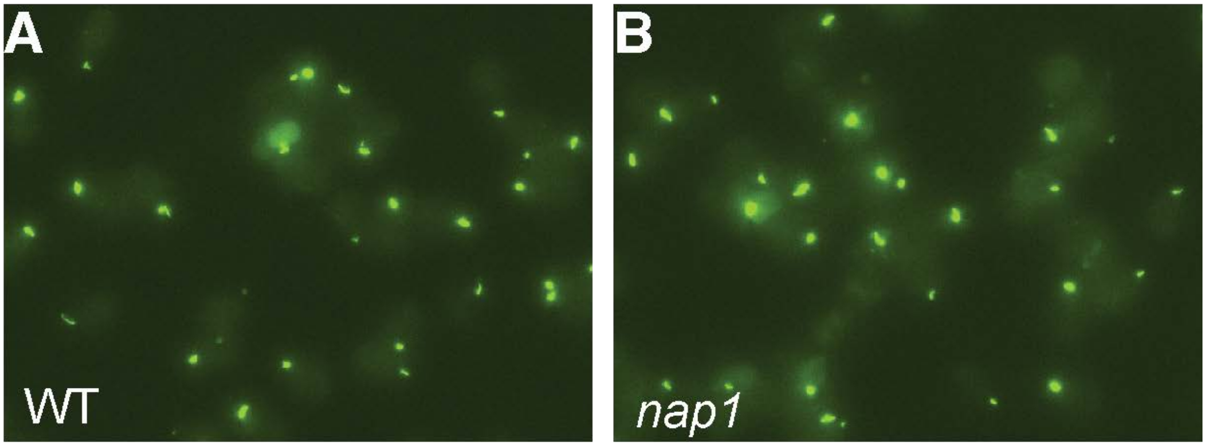


**Supplementary Figure S4: Mutants of NAP1 have unaffected fertilization tubule formation.**

**(A-B)** Wild-type and *nap1-1* cells stained with phalloidin-Alexa Fluor 488 to label actin-rich fertilization tubules.

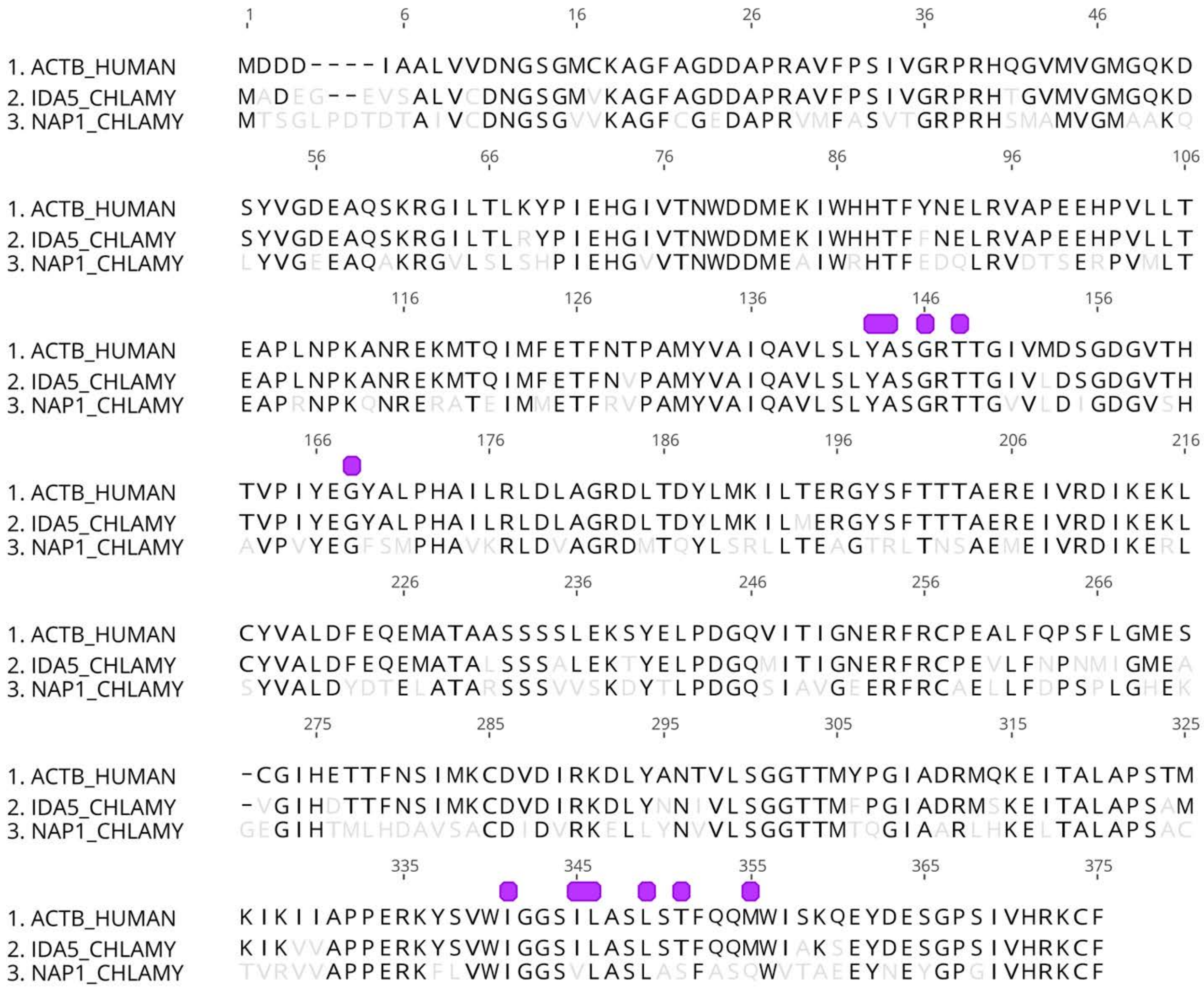


**Supplementary Figure S5: ClustalW alignment of human actin with *Chlamydomonas* actins.** The actin-binding FH2 domain of formin interacts with a hydrophobic cleft between actin subdomains 1 and 3 lined by the residues indicated in purple. Human actin is aligned with conventional (IDA5) and unconventional (NAP1) *Chlamydomonas* actin. Non-conserved residues are in light gray.

**Supplemental Movie Figure Legends:**

**Movie 1: PRF1 inhibits Arp2/3 complex-mediated actin assembly, related to Figure 6.** TIRF microscopy bead assays. Beads coated with fission yeast Wsp1 were incubated with a series of components (listed in top left). Wsp1 bead is incubated with 1.5 μM actin (10% Alexa-488 labeled) and 30 nM Arp2/3 complex (Actin, Arp2/3) followed by flowing in a mixture of the same concentrations of actin, Arp2/3 complex, and 2.5 μM PRF1 (+PRF1). The bright flash in each movie indicates photobleaching, which allows visualization of exclusively new actin assembly. Scale bar, 5 μm. Time in sec.

**Movie 2: PRF1 promotes FOR1-mediated actin assembly, related to Figure 6.** TIRF microscopy bead assays. Beads coated with FOR1(3P,FH2) were incubated with a series of components (listed in top left of each movie). FOR1 bead is incubated with 1.5 μM actin (10% Alexa-488 labeled) (Actin), followed by flowing in a mixture of actin and 2.5 μM PRF1 (+PRF1). The bright flash indicates photobleaching, which allows visualization of exclusively new actin assembly. Scale bar, 5 μm. Time in sec.
